# Transient immune landscape remodelling shapes CD8 T-cell priming during infection

**DOI:** 10.64898/2026.03.18.712682

**Authors:** Ana Laura Chiodetti, Han Cai, Andrew Muir, Gracie J. Mead, Mariana Borsa, Arianne C. Richard, Audrey Gérard

## Abstract

Immune responses to intracellular pathogens require the generation of a CD8 T-cell response comprised of clones with a broad range of T Cell Receptor (TCR) affinities for their cognate peptide. While high-affinity clones clear pathogens during initial infection, low-affinity T-cells are critical to protect against subsequent encounters with mutated pathogens. Low-affinity T-cells are recruited to an immune response despite known mechanisms that limit their expansion and priming. As such, the mechanism safeguarding the recruitment of low-affinity T-cell clones during infection remains unclear. Here, we show that at the start of an infection, the spatial immune architecture is transiently reorganised to initiate broad T-cell priming. In particular, Tregs become segregated from the initial priming site, resulting in a spatio-temporal window during which pathogen-specific T-cell priming is protected from Treg inhibition. Priming induces the expression of CXCR3 in a TCR affinity-dependent manner, which conditions the retention of activated T-cells in the priming region. Inability to express CXCR3 leads to T-cell relocation towards Treg-rich areas, where their expansion is limited by the lack of the costimulatory molecule CD70. Our study redefines the role of Treg during infection and provides a mechanism to explain how the priming and breadth of the CD8 T-cell response is protected during infection.

## Introduction

CD8 T-cell priming occurs in secondary lymphoid organs (SLOs), where their T-Cell Receptor (TCR) engages its cognate peptide coupled to MHC-I (pMHC-I) on antigen-presenting cells (APCs). This is concomitant with the engagement of co-stimulatory molecules, necessary to trigger CD8 T-cell proliferation^1^, and inflammatory cues that skew their differentiation^2^. Each T-cell recruited to an immune response expresses a distinct TCR that recognises pathogen-derived pMHC-I with an affinity that can vary over multiple orders of magnitude^3^. Overall, the average avidity of the T-cell response has to be high enough to control pathogens. But low-affinity T-cells also need to be actively recruited. In fact, low-affinity T-cells are the ones responding to mutated pathogens during secondary infections, driving heterosubtypic immunity^4^. Recruiting a large breadth of TCR affinities is therefore critical^5,6^.

While recruitment of low-affinity T-cells during infection is well-documented^5–7^, what safeguards their recruitment remains unclear. Indeed, multiple mechanisms are in place to increase the threshold of T-cell priming and promote the preferential expansion of high-affinity T-cells. High-affinity T-cells have increased expression of Myc^8,9^ and IRF4^10^, which leads to higher intrinsic capacity to expand^11^. In addition, extrinsic mechanisms also increase the affinity and magnitude of the T-cell response. Those include the presence of Tregs which have been suggested to further restrict low-affinity T-cell priming and overall T-cell expansion^12–16^ through multiple mechanisms including IL2 sequestration^15^, CD73-derived adenosine production^14^, TGFß secretion^14^, inhibition of Dendritic Cell (DC) activation^17–20^ and restriction of chemotactic cues^12^. But despite this, low-affinity T-cells get recruited during infection.

How can the immune system simultaneously increase the TCR affinity threshold while safeguarding the recruitment of low-affinity T-cells? One hypothesis is that the control of affinity threshold and priming of low-affinity T-cells rely on distinct signals present in dedicated spatiotemporal niches. We and others have pioneered the study of spatiotemporal T-cell priming and differentiation^21^, demonstrating that the generation of distinct cell-cell communication platforms and microenvironments guides T-cell differentiation. Initial priming of CD8 and CD4 T-cells happens in distinct niches^22,23^, followed by migration to a common site, allowing CD8 T-cells to get CD4 T-cell help, fostering memory generation^22,23^. In addition, priming in a specific innate cell environment skews the fate of CD8 T-cells during type-I inflammation^24^. Finally, we reported that T-cells are primed separately in SLOs and then aggregate with each other, influencing their fate through integrin-mediated interactions^25,26^. Whether similar niche formation is required to enable low-affinity priming is unknown.

Here, we used a combination of imaging, photoactivation, and RNA-sequencing to identify the critical spatio-temporal features required for priming a broad CD8 T-cell response during infection. Using Listeria monocytogenes infection, we show that during the inception of the CD8 T-cell response, Tregs are segregated from the initial priming site. As a result, there is a temporal window during which pathogen-specific T-cell priming is protected from Treg inhibition. The initial priming site is rich in innate cells such as NK cells and monocytes and inflammatory cytokines, and supports efficient T-cell priming and expansion through CD70-expressing dendritic cells. Priming induces CD8 T-cell expression of CXCR3 in a TCR affinity-dependent manner, and CXCR3-dependent retention of primed T-cells in innate-infiltrated regions. Priming in innate-infiltrated regions is required for initiating broad T-cell priming, as CXCR3 inhibition leads to the enrichment of primed T-cells in Treg-rich areas of T-cell zones, where their expansion is compromised. Our study redefines the role of Tregs during infection and provides a mechanism to explain how the breadth of an immune response can be protected.

## Results

### High- and low-affinity CD8 T-cells exhibit similar spatio-temporal priming pattern

To investigate whether the spatio-temporal behaviour of CD8 T-cells was affected by TCR affinity, we first characterised the different niches formed during infection in the spleen, using *Listeria monocytogenes* (LM) as a common model to study CD8 T-cell responses to blood-borne pathogens. The distribution of immune cells was analysed by confocal microscopy in the spleens of mice infected by LM over time. The marginal zone (MZ) macrophage marker CD169 delineated the red pulp (RP) from the white pulp (WP), and CD11b and NKp46 were used to track monocytes and natural killer (NK) cells, respectively. Consistent with previous reports on other infection models^27–30^, we observed a transient spatial reorganisation of immune cells within the WP. NK cells moved from the RP and partially infiltrated the WP within the first 12-18 hours following infection (*Fig.1A*). Recruitment of monocytes to the WP also occurred and followed the same kinetics as NK cells (*Fig.1A*). Innate infiltration increased up to 24 hours, when innate cells formed dense clusters (*Fig.1A*), and inflammatory cells left the WP by Day 2 to 3 (*Fig.S1A*), highlighting the transitory nature of innate cell infiltration to the WP.

**Figure 1.**
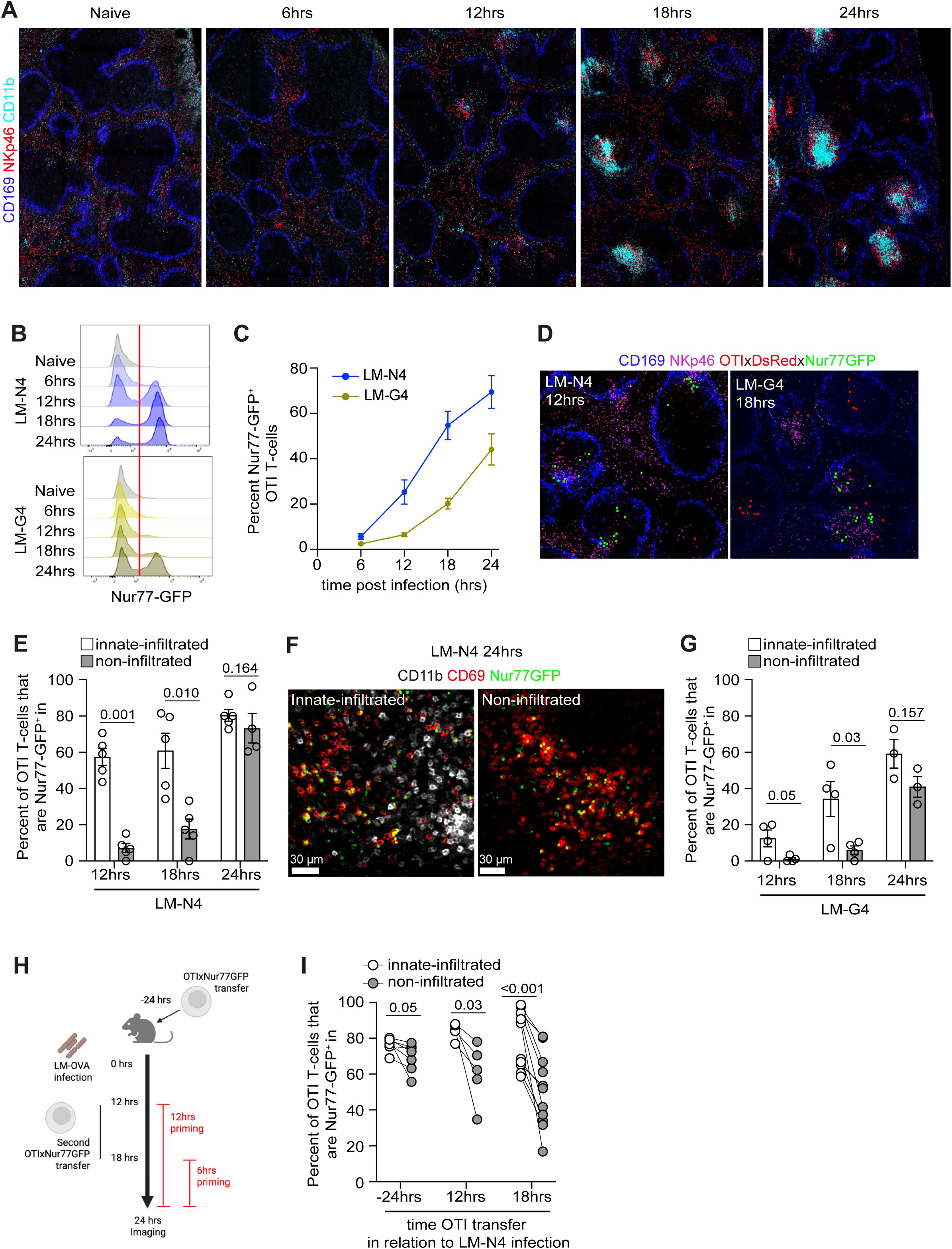
CD8 T-cell priming is spatiotemporally linked to inflammatory innate cell infiltration following Listeria monocytogenes infection. **(A-G)** WT mice were adoptively transferred with OTIxhCD2-DsRedxNur77-GFP **(A-E, G)** or OTIxNur77-GFP **(F)** T-cells and infected with LM-N4 or LM-G4. Spleens were harvested at the indicated time points and processed for flow cytometry and immunofluorescence imaging. **A-** Immunofluorescence images of spleen sections. CD169 (blue), CD11b (cyan), NKp46 (red). **B-** Representative histograms showing Nur77-GFP expression in OTI T-cells over time. **C-** Percentages of Nur77-GFP+ OTI T-cells over time (n=4 for 6hr, n=12 for 12hr, n=16 for 18hr, n=17 for 24hr). **D-** Immunofluorescence images showing the positioning of Nur77-GFP- OTI T-cells (red dots) and Nur77-GFP+ OTI T-cells (green dots) in relation to NK cells (magenta) in spleens infected with LM-N4 and LM-G4 12hr and 18hr prior, respectively. **E-**Percentages of Nur77-GFP+ OTI T-cells located within innate-infiltrated and non-infiltrated regions over time following LM-N4 infection (n=9-10 per timepoint). **F-** Representative images showing the positioning of Nur77-GFP+ OTI T-cells (green), CD69 expression (red) and CD11b+ cells (white) in spleens infected with LM-OVA for 24hrs. **G-** Percentages of Nur77-GFP+ cells among OTI T-cells located within innate-infiltrated and non-infiltrated regions on stained spleen sections at indicated timepoints following LM-G4 infection (n=6-8 per timepoint). **(H-I)** WT mice infected with LM-N4 were adoptively transferred with OTI-RFPxNur77-GFP T-cells or Far-red dye-labelled OT1xNur77-GFP at indicated time points. Spleens were harvested 24hr post-infection **(H)**. **I-** Percentages of Nur77-GFP+ OTI T-cells located within innate-infiltrated and non-infiltrated regions. Multiple paired t-tests with multiple comparisons correction using Holm-Šídák method **(E, G)**, Paired t-tests (**I**). Innate-infiltrated regions are defined as have ≥ 1 NK cell per 10^3^μm^2^ **(E,G,I)**.

To characterise how innate infiltration integrates with T-cell priming, we transferred OVA-specific CD8 T-cells (OTI T-cells) into mice that were then infected with LM. LM were expressing the OVA peptide SIINFEKL (N4) (LM-N4) or the altered OVA peptide SIIGFEKL (G4) (LM-G4), recognised by the OTI TCR with high- or low-affinity, respectively^31,32^. These OTI T-cells also expressed the Nur77-GFP reporter (OTIxNur77-GFP), whereby TCR triggering induces GFP expression^33^. GFP expression was detected in OTI T-cells 12 hours following infection with LM-N4, and most cells were activated by 24hrs (*Fig.1B-C, S1B*). When mice were infected with LM-G4, although the percentage of activated OTI T-cells was lower, the kinetics were similar (*Fig.1B-C*), showing that initial CD8 T-cell priming occurs within the first 24hrs following LM infection. Similar data were observed when we assessed the activation markers CD69 and CD25 (*Fig.S1C*). This demonstrates that the kinetics of T-cell priming paralleled that of innate cell infiltration. In fact, T-cell priming, visualised *in situ* by Nur77-GFP expression, occurred where innate cells infiltrated (*Fig.1D*), suggesting that antigen was mostly presented in innate-infiltrated regions. However, quantification of Nur77-GFP positive OTI T-cells over time demonstrated that, while they were first detected in innate-infiltrated regions, they were also found in non-infiltrated regions by 24hrs (*Fig.1E-F*). The frequency of T-cells that were positive for GFP was similar between regions 24hrs post-infection, suggesting similar antigen availability at that time. Here again, the localisation pattern of primed T-cells was not driven by TCR affinity, as OTI T-cell priming following LM-G4 infection showed the same dynamics over time (*Fig.1G*).

While we demonstrated that primed CD8 T-cells were initially located in innate-infiltrated regions, and subsequentially in both innate-infiltrated and non-infiltrated regions, it was not clear whether (i) antigen had spread to non-infiltrated regions and T-cells randomly found their antigen in either region at later times, or (ii) T-cells were primed in innate-infiltrated regions and moved to non-infiltrated regions when they resumed migration. To differentiate between both hypothesises, we infected mice with LM-N4 and injected OTIxNur77-GFP T-cells at later time points, 12hrs or 18hrs after infection (*Fig.1H*). The frequency of OTI T-cells that expressed GFP in each region 24hrs post infection was then quantified by imaging. If equal priming in both regions at 24hrs was due to similar antigen availability, we would expect T-cells from delayed transfer to be primed equivalently in either region. However, if CD8 T-cells were first primed in innate-infiltrated regions and then moved in non-infiltrated regions, we would expect OTI T-cells that were transferred at later time points to preferentially display signs of priming in innate-infiltrated regions. OTI T-cells transferred 12hrs after infection preferentially expressed GFP in innate-infiltrated regions compared to non-infiltrated regions. This was even more accentuated when T-cells were transferred 18hrs after infection (*Fig.1I*), when they only had the opportunity to get primed for 6hrs. This is consistent with a scenario where most CD8 T-cells were primed in innate-infiltrated regions and a fraction then moved to non-infiltrated regions at later times.

Altogether, our data indicate that most T-cells are initially primed in innate-infiltrated regions that are rich in NK cells and monocytes, and at the peak of priming, a fraction of activated T-cells migrate to non-infiltrated regions. T-cells follow this priming pattern regardless of TCR affinity.

### CXCR3 expression conditions CD8 T-cell retention in innate-infiltrated regions in a TCR affinity-dependent manner, required for their expansion

We then set out to understand what signals govern CD8 T-cell localisation in innate-infiltrated or non-infiltrated regions. Innate-infiltrated regions are characterised by the presence of NK cells, which are known to produce IFNγ early after infection^26,34,35^. Indeed, the expression of IFNγ was restricted to innate-infiltrated regions (*Fig.S2A*), as reported in multiple immunisation models^27,35–37^. IFNγ is known to induce the expression of the chemokines CXCL9 and CXCL10, and their receptor CXCR3 is upregulated in CD8 T-cells following TCR stimulation^38,39^. We therefore hypothesised that CXCL9/10 was produced in innate-infiltrated regions and promoted T-cell retention. Analysis of CXCL9 distribution 24hrs after LM infection revealed that CXCL9 expression was indeed enriched in innate-infiltrated regions (*Fig.2A-B*).

**Figure 2.**
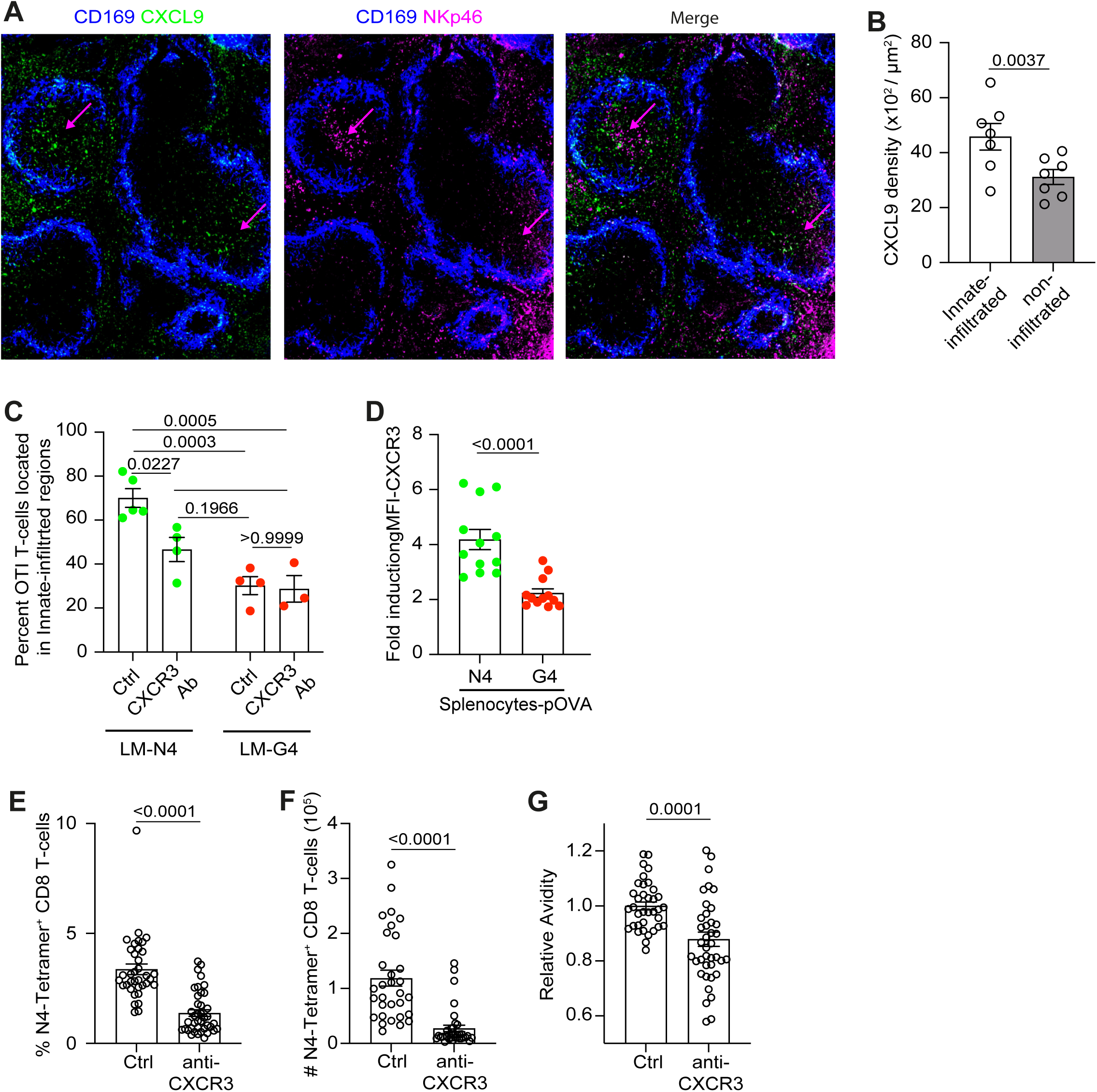
CXCR3 expression on high-affinity CD8 T-cells allows their retention in innate-infiltrated regions and promotes expansion. **(A-B)** Spleen sections from mice infected with LM-OVA for 24hrs were stained for CD169 (blue), CXCL9 (green), NKp46 (magenta). **A-**Representative images. **B-** Average CXCL9 signal intensities within innate-infiltrated (NKp46+) and non-infiltrated (NKp46-) regions (n=6). **C-** Mice were adoptively transferred with OTI-hCD2-DsRed cells, infected with LM-N4 or LM-G4 and treated with either anti-CXCR3 or isotype control antibodies 6hr prior to infection. Spleen sections were stained for CD169 (blue) and NKp46 (magenta) (n=3 per group) after 24hrs. Percentages of OTI T-cells located within innate-infiltrated regions. **D-** Isolated OTI T-cells were cocultured with IFNy-primed WT splenocytes in the presence of N4 or G4 peptides. CXCR3 expression was examined 24hr later by flow cytometry. Data is expressed as CXCR3 fold-induction on CD69+ CD8 OTI T-cells relative to unstimulated condition (n=4). (**E-G)** Mice were treated with either isotype control or anti-CXCR3 antibodies and infected with LM-OVA 6hr later. Spleens were harvested 8 days post-infection and analysed by flow cytometry (n=36-40 per group). Data shows frequencies **(E)** and absolute numbers **(F)** of N4-tetramer+ CD8 T-cells. **G-** Graph shows relative avidity, calculated by dividing the N4-tetramer gMFI to CD3 gMFI, of N4-tetramer+ CD8+ T cells, normalised to isotype control. Ratio paired t-test **(B)**, Two-way ANOVA with Turkey’s multiple comparison test **(C)**, Welch’s t-tests **(D-G)**. Data show mean ± SEM.

This data therefore suggests that CXCR3 and its ligands might contribute to CD8 T-cell retention in innate-infiltrated regions after their initial priming. To test this, we inhibited CXCR3 using blocking antibodies and analysed OTI T-cell location 24hrs after infection with LM-N4. As a control, we observed that blocking CXCR3 did not affect the size and number of innate-infiltrated regions (*Fig.S2B*). Blocking CXCR3 resulted in OTI T-cell redistribution from innate-infiltrated regions to non-infiltrated areas (*Fig.2C, S2C*), confirming that CXCR3 is important for the retention of CD8 T-cells in innate-infiltrated regions. However, OTI T-cells preferentially accumulated in non-infiltrated regions when they were primed with the low-affinity peptide G4 and blocking CXCR3 had little effect on their location (*Fig.2C, S2C*). This suggested that CXCR3 was not promoting low-affinity CD8 T-cell retention in innate-infiltrated regions. Therefore, we concluded that CXCR3 expression, likely on CD8 T-cells, conditions T-cell retention in innate-infiltrated regions in a TCR affinity-dependent manner. Consistent with this, while CXCR3 was upregulated in CD8 T-cells 24hrs after TCR priming following *in vitro* activation (*Fig.2D*), the extent of CXCR3 expression was TCR affinity dependent^15,38^, as priming with the low-affinity peptide G4 results in mild upregulation of CXCR3 by OTI T-cells compared to priming with the high-affinity peptide N4 (*Fig.2D*).

To explore the functional relevance of CD8 T-cell retention in innate-infiltrated regions, we inhibited CXCR3 specifically during priming. To do so, WT mice were infected with LM-OVA and treated with a CXCR3-blocking antibody once 6hrs before infection. Splenocytes were isolated at the peak of the infection and OVA-specific T-cells were quantified by N4-tetramer staining (*Fig.S2D*). Blocking CXCR3 during priming was sufficient to limit the number and percentage of OVA-specific T-cells (*Fig.2E-F*), suggesting that retention in innate-infiltrated regions was critical to potentiate CD8 T-cell expansion. Consistent with the fact that CXCR3 blocking affected high-affinity T-cell location, we found that blocking CXCR3 during priming lowered the avidity of the T-cell response (*Fig.2G*), assessed by normalising tetramer staining to CD3 staining^35^. Interestingly, early CXCR3 blockade only had subtle effect on T-cell phenotype at the peak of the response (*Fig.S2E*), suggesting that CXCR3-dependent retention in innate-infiltrated regions is important for T-cell expansion and avidity rather than differentiation. Finally, as CXCR3 is also important at later stages for T-cell differentiation^40–43^, we blocked CXCR3 between day 3 and day 7 post-infection and analysed T-cell expansion and avidity at the peak of the effector response. While CXCR3 was still important at later time points for T-cell expansion (*Fig.S2F*), most likely due to the requirement of CXCR3 expression for effector differentiation^40–43^, blocking CXCR3 between day 3 and day 7 post-infection did not affect the avidity of the effector response (*Fig.S2G*). This confirms that CD8 T-cell retention in innate-infiltrated regions is important for their expansion and controls the avidity of the response.

Overall, we concluded that CD8 T-cells, after being primed in innate-infiltrated regions, express CXCR3 in a TCR affinity-dependent manner. If they fail to do so, they are more likely to migrate to non-infiltrated regions, which restricts their expansion.

### CD8 T-cell dynamics suggest distinct priming cues in innate-infiltrated and non-infiltrated region

Our data so far demonstrate that CD8 T-cell localisation in innate-infiltrated regions is important for expansion, but T-cells also displayed signs of TCR activation, such as clustering and expression of CD69, in non-infiltrated regions (*Fig.S3A*). They also expressed the reporter Nur77-GFP in both regions, demonstrating that they have recently seen their antigen (*Fig.1E-G*). This suggested that CD8 T-cells also encountered antigen in non-infiltrated regions, but the quality of TCR activation or quantity of antigen there might be sub-optimal.

To address whether CD8 T-cells indeed encountered antigen in non-infiltrated areas, we turned to 2-photon microscopy to characterise the dynamics of high- and low-affinity T-cells in each region. We co-transferred OTI T-cells with OT3 T-cells, a CD8 T-cell clone that expresses a TCR recognising the N4 peptide with 50-fold decreased functional avidity compared to the OTI TCR^44^. Both clones were tagged with a fluorescent protein to track them and compare their behaviours in the same mouse. Mice were infected with LM-OVA and spleens were collected after 24hrs for 2-photon imaging. CD11b or Nkp46 staining was used to distinguish innate-infiltrated from non-infiltrated regions. OT3 T-cells preferentially accumulated in non-infiltrated regions (*Fig.S3B*), as seen for OTI T-cells primed with low-affinity (LM-G4, *Fig.S2C*), reinforcing the impact of TCR affinity on localisation and validating the use of OT3 T-cells as a low-affinity T-cell clone.

Active TCR priming is characterised by T-cell clustering and arrest^45–47^. We found clusters between OTI and OT3 T-cells in innate-infiltrated and non-infiltrated regions (*Fig.3A-B*), and OTI and OT3 T-cell displacement were decreased in both regions compared to naïve animals (*Fig.3C-D*). They also displayed lower speed (*Fig.3E*) and increased arrest (*Fig.3F*), overall confirming that T-cells were seeing their antigen in both microenvironments (*Video S1-3*). T-cells present in innate-infiltrated niches exhibited decreased displacement, lower speed and higher arrest coefficient than the ones present in non-infiltrated regions (*Fig.3C-F*), consistent with qualitative and/or quantitative differences in priming. Surprisingly, we observed only marginal difference in speed (*Fig.3E*) and arrest (*Fig.3F*) between OTI and OT3 T-cells within each niche. This indicates that, while TCR affinity causes preferential regional localisation, upon entering a region, TCR affinity does not affect T-cell motility. Other studies described differences in high-and low-affinity T-cell motility following DC or peptide immunisation^38,48,49^, suggesting that T-cell behaviour in the context of immunisation and infection is differentially regulated. Finally, the same dynamics were observed for OTI and OT3 T-cells when transferred into different recipients, suggesting that clonal competition has no significant impact on their dynamics (*Fig.S3C-F*).

**Figure 3.**
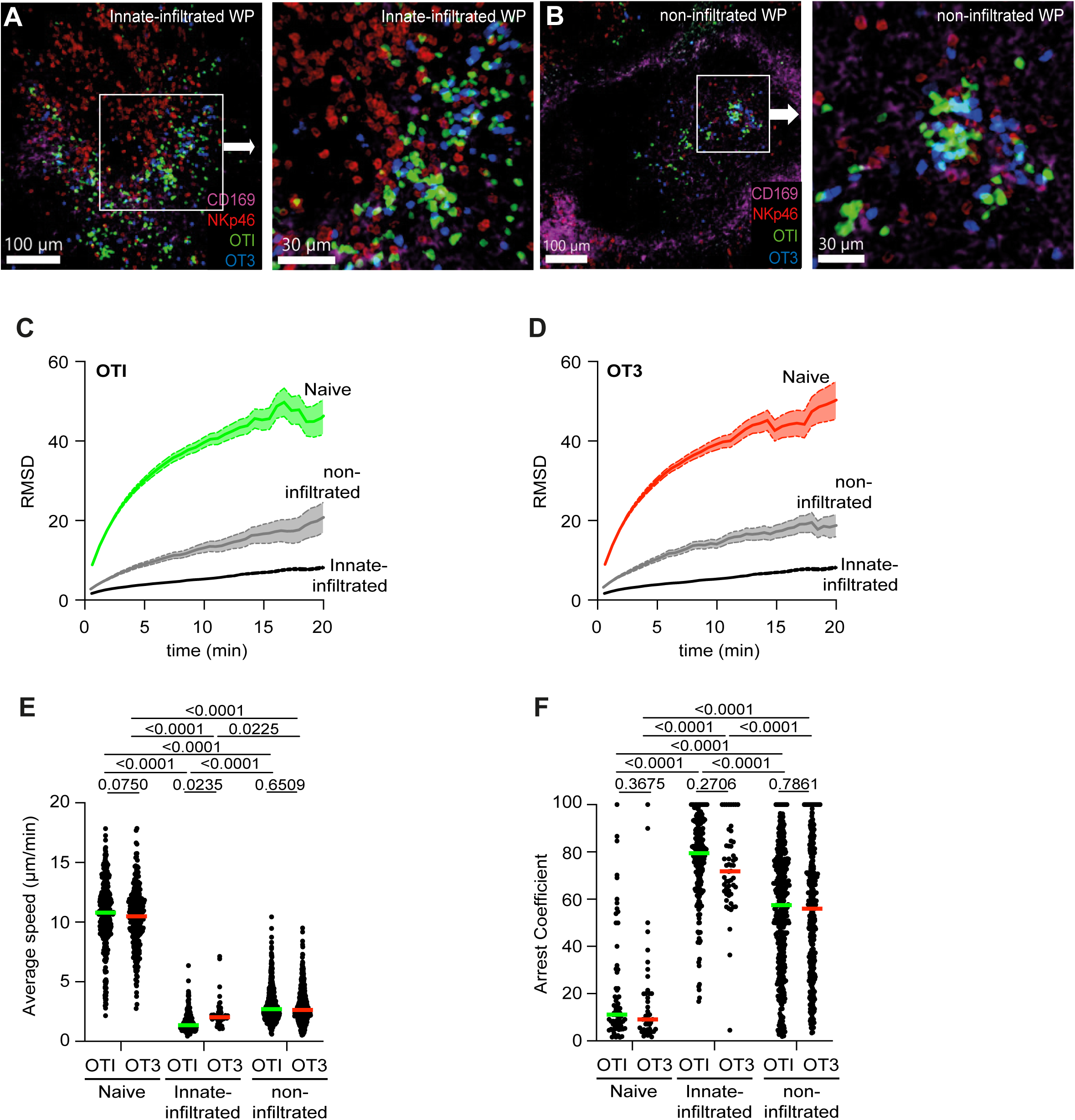
CD8 T-cells encounter antigens in both innate-infiltrated and non-infiltrated regions, regardless of TCR affinity. Mice were adoptively transferred with admixed OTIxGFP and OT3xhCD2-DsRed cells and infected with LM-OVA. After 24hrs, live spleen sections were labelled with NKp46 **(A-B)** or CD11b **(C-F)** and CD169 and imaged by 2-photon microscopy. Data is representative of 3 independent experiments with at least 3 spleens per condition. (**A-B**) Image of innate-infiltrated (**A**) and non-infiltrated (**B**) regions in a WP showing clustering of OTI (green) and OT3 (blue) T-cells. Left: whole WP (scale bar = 100um); Right: zoom (scale bar = 30um). (**C-D**) Graphs show the Root Mean Square Displacement (RMSD) over time of OTI (**C**) and OT3 (**D**) cells in naive (OTI:green/OT3:red), innate-infiltrated (grey) and non-infiltrated (black) regions. (**E-F**) Average speed (**E**) and arrest coefficient (**F**) of OTI (green) and OT3 (red) T-cells in naive, innate-infiltrated or non-infiltrated regions. Each dot is a cell, bar indicates Median.

This data overall confirms that high- and low-affinity CD8 T-cells present in non-infiltrated regions are seeing antigen but suggests a lower quality of priming in non-infiltrated regions. This is consistent with decreased CD8 T-cell expansion observed when T-cells are enriched in non-infiltrated regions following CXCR3 blockade.

### Expression of CD70 is restricted to innate-infiltrated regions and required for T-cell expansion

Based on the different CD8 T-cell dynamics between innate-infiltrated and non-infiltrated regions, we investigated the type and quality of priming in each region. We first analysed the location of dendritic cell (DC) subsets 24hrs after LM-OVA infection, as DC subsets have different capacities to prime CD8 T-cells^50^. cDC1, a subset of DCs specialised in cross-presentation, was identified by XCR1 staining. The density of cDC1 was higher in non-infiltrated regions (*Fig.4A, S4A-B*). Inversely, cDC2, a subset associated with CD4 T-cell priming and identified by SIRP1a or 33D1 staining, showed higher density in innate-infiltrated regions (*Fig.4B, S4A-B*). Importantly, while cDC1s were sparse in innate-infiltrated regions, they were still present and supported OTI T-cell clustering (*Fig.4C, S4C*), suggesting they play a significant role in CD8 T-cell priming in innate-infiltrated regions.

**Figure 4.**
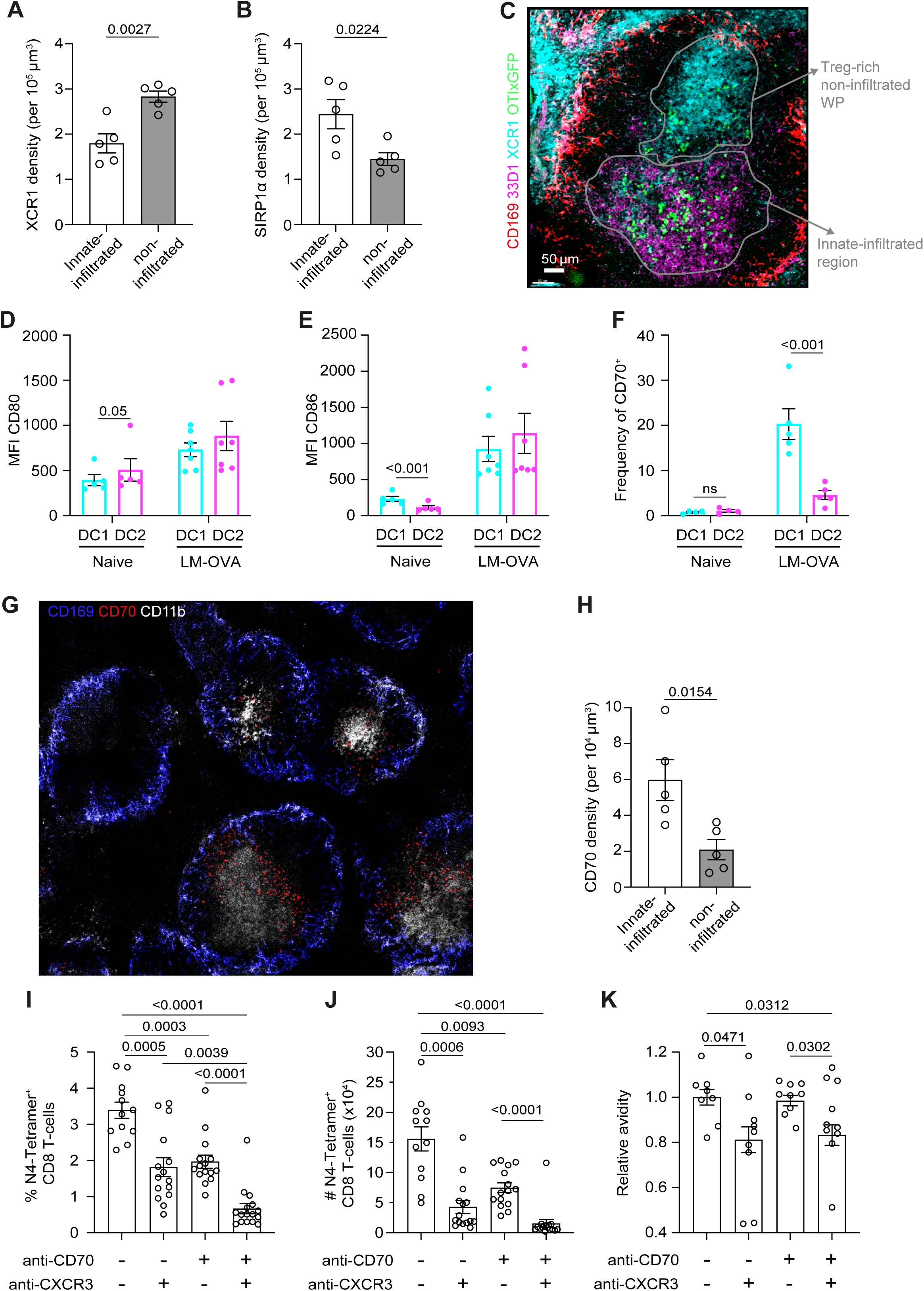
CD70 is enriched within innate-infiltrated regions and supports CD8 T-cell expansion. **(A-B)** Graphs show XCR1 **(A)** and SIRPa **(B)** signal density within innate-infiltrated and non-infiltrated regions on stained spleen sections harvested 24hrs after LM-OVA infection (n=5). **C-**Mice were adoptively transferred with OTIxGFP T-cells and infected with LM-OVA. After 24hrs, spleen sections were stained for CD169 (red), 33D1 (magenta), XCR1 (cyan), OTI (green). Representative image. **(D-F)** Mice were infected with LM-OVA for 24hrs and splenocytes were analysed by flow cytometry. Graphs show CD80 MFI (**D**, n=5-7), CD86 MFI (**E**, n=5-7), and frequency of CD70+ cDC1 and cDC2 (**F**, n=4-5). **(G-H)** Spleen sections from mice infected with LM-OVA for 24hrs were stained for CD169 (blue), CD70 (red), CD11b (white). **G-** Representative image. **H-** Graph shows CD70 signal intensity within innate-infiltrated and non-infiltrated regions in WPs (n=5). **I-K** Mice were infected with LM-OVA and treated with anti-CXCR3 or anti-CD70 antibodies as indicated 6hrs before infection. Graphs show frequencies **(I)**, absolute numbers **(J)** and avidity **(K)** of N4-tetramer+ CD8 T-cells 8 days post-infection (n=12-17 per group). Ratio paired t-tests **(A-B, D-F, H)**, Welch and Brown-Forsythe one-way ANOVA with Dunnett’s multiple comparisons test **(I-K)**.

The expression of costimulatory molecules is increased during infection^51^, and some are regulated by inflammation^52^. Given the generation of specific innate-infiltrated niches characterised by inflammatory mediators during T-cell priming, we wondered whether costimulatory ligand expression might differ between regions. We first analysed the expression of CD80 and CD86. Following LM-OVA infection, both costimulatory molecules were up-regulated by cDC1 and cDC2 (*Fig.4D-E* and *S4D-E*), which implies that their expression was not restricted to specific regions. We then analysed the expression of CD70, the ligand of TNF family receptor CD27, as it is upregulated during inflammatory conditions^53,54^ and important for low-affinity T-cell priming^55^. Using flow cytometry, we found that CD70 expression was restricted to a fraction of XCR1^+^ cDC1 (*Fig.4F and S4D-E*). Imaging spleen sections 24hrs following LM-OVA infection demonstrated that CD70 density was higher in innate-infiltrated compared to non-infiltrated regions (*Fig.4G-H*), suggesting that the few cDC1 that populated innate-infiltrated regions were providing CD70-dependent costimulation. Altogether, this data indicated that CD70-expressing cDC1 present in innate-infiltrated regions were key to priming CD8 T-cells. To explore this, we blocked CD70 specifically during priming using a single dose of CD70 blocking antibody 6hrs before LM-OVA infection and analysed the endogenous response against OVA using N4-tetramers. Blocking CD70 limited OVA-specific CD8 T-cell expansion (*Fig.4I-J*). Importantly, blocking both CXCR3 and CD70 had a combinatorial effect and fully abrogated OVA-specific CD8 T-cell expansion, demonstrating that these two signals are critical for CD8 T-cell expansion (*Fig.4I-J*). In addition, while blocking CXCR3 decreased the avidity of the CD8 T-cell response, blocking CD70 alone had no effect on avidity, indicating that both high- and low-affinity CD8 T-cells relied on CD70 for their priming and subsequent expansion.

Altogether, our data demonstrate that CD70 expression is spatially restricted to areas of innate infiltration and is required for efficient CD8 T-cell expansion, explaining why T-cells retained in innate-infiltrated regions expand more.

### Differences in CD8 T-cell priming in innate-infiltrated and non-infiltrated niches

To confirm that TCR priming in non-infiltrated regions was not as effective as priming in innate-infiltrated regions, we compared the transcriptome of CD8 T-cells present in innate-infiltrated and non-infiltrated regions using a strategy based on photoactivatable-GFP (PA-GFP) (*Fig.5A*). PA-GFP is a variant of GFP that is not fluorescent at steady-state but becomes fluorescent when activated using 2-photon excitation^56^. PA-GFP mice were crossed to hCD2-DsRed mice to label T-cells with DsRed, highlighting WPs. Mice were infected with LM-OVA. After 24 hours, spleens were explanted and live spleen sections were stained for CD11b and Ly6c to delineate innate-infiltrated regions (*Fig.5B*). Cells within innate-infiltrated or non-infiltrated regions were then labelled by photoactivating PA-GFP (*Fig.5C*). GFP^+^ CD8 T-cells were sorted and subjected to bulk RNAseq. When compared to CD8 T-cells from naïve spleens, CD8 T-cells from innate-infiltrated regions of infected spleens exhibited more differential gene expression than those from non-infiltrated regions (*Fig.5D*). Running gene set enrichment analysis on genes differentially expressed in CD8 T-cells from innate-infiltrated versus non-infiltrated regions of infected spleens (*Fig.S5A, Dataset 1*) revealed that cells in innate-infiltrated regions preferentially expressed genes related to the adaptive immune response and regulation of protein modification and metabolism (*Fig.5E*, *Dataset 2*). Those pathways are consistent with T-cell recruitment to an immune response and expansion. Interestingly, only a few genes were differentially expressed between naïve CD8 T-cells and those isolated from non-infiltrated regions (*Fig.5D*). This confirms that the quality of TCR priming is different between innate-infiltrated and non-infiltrated regions.

**Figure 5.**
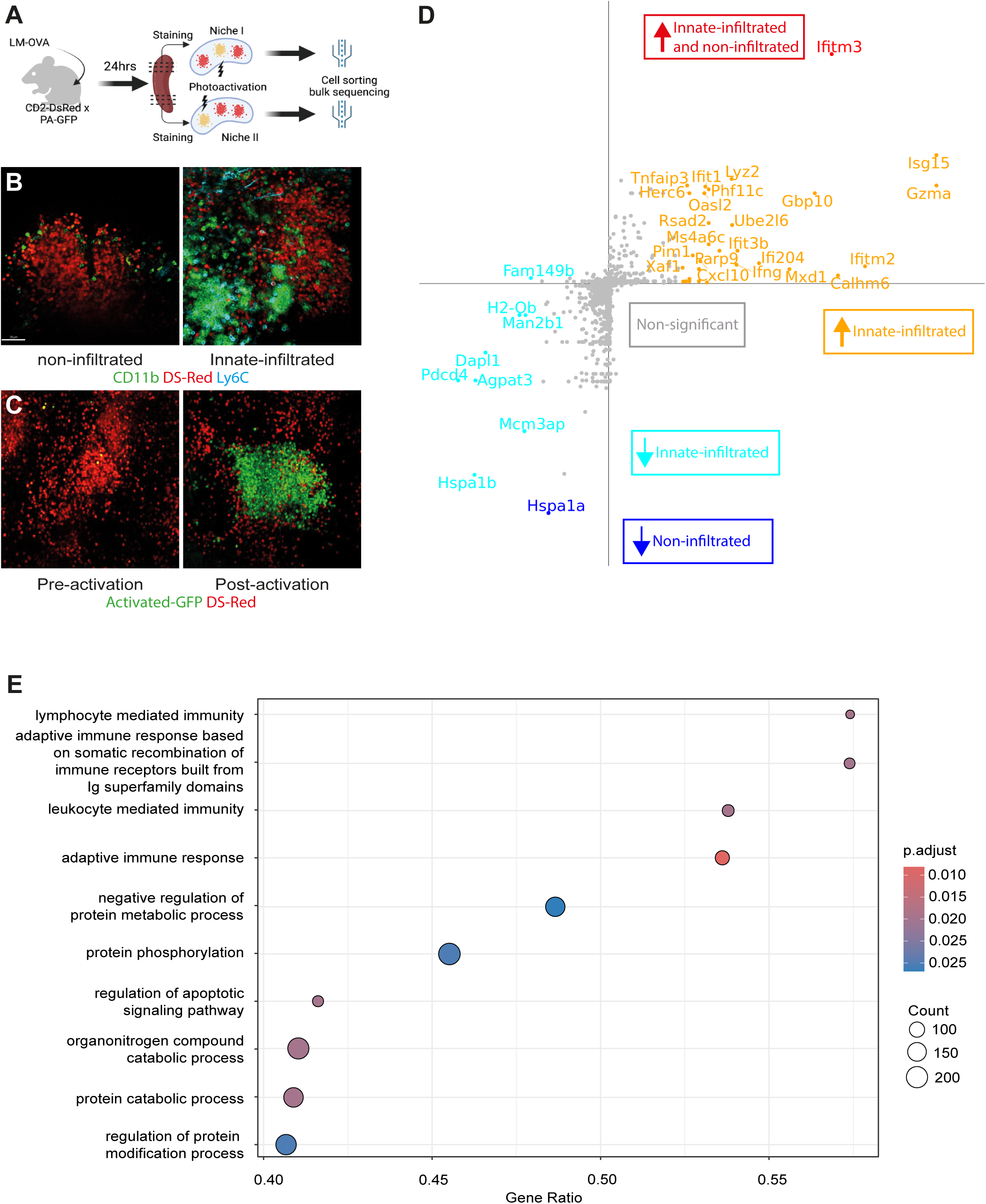
Transcriptomics analysis of CD8 T-cells in innate-infiltrated and non-infiltrated regions. hCD2-DsRedxPA-GFP mice were infected with LM-OVA. After 24hrs, spleens were explanted, sectioned and stained for CD11b and Ly6C. Innate-infiltrated and non-infiltrated regions were highlighted by 2-photon light to activate PA-GFP. PA-GFP+ cells were then sorted and subjected to bulk-RNAseq. **A-** Experimental set-up. **B-** Example of non-infiltrated (left) and innate-infiltrated (right) regions. **C-** Example of DsRed+ cells before (left) and after (right) photoconversion. **D-** Differential expression analysis displaying genes significantly up-regulated (orange) and down-regulated (cyan) in innate-infiltrated niches, downregulated in non-infiltrated niches (blue) and up-regulated in innate-infiltrated and non-infiltrated niches (red) compared to naïve. **E-** Gene set enrichment analysis of T-cells from innate-infiltrated versus non-infiltrated niches.

These data provided further evidence that the location of CD8 T-cells influences the type of priming they experience. We aimed to confirm this by analysing scRNAseq data and asked whether we could identify T-cells exhibiting the gene signature elicited in naïve mice, or innate-infiltrated and non-infiltrated regions of infected mice (*Dataset 3-4*). We selected scRNAseq data from CD8 T-cells isolated from naïve spleens or spleens 24hrs after LM-OVA infection^35^, excluding memory T-cells. We used SingleR to label cells as belonging to the naïve pool, innate-infiltrated or non-infiltrated regions, by using our sorted bulk RNAseq data as an atlas. Clustering delineated 5 clusters (C0 to C4), with C0 and C1 present in naïve and infected mice and C2-4 enriched in Listeria-infected mice (*Fig.S5B*). Interestingly, cells labelled as naïve were not found in LM-OVA-infected mice, suggesting that infection had a widespread effect on T-cells (*Fig.S5B-C*). In LM-OVA-infected spleens, about half of the cells were labelled as Innate-infiltrated (*Fig.S5B-C*), which were enriched in C2-4 (*Fig.S5D*). Further Gene Ontology enrichment analysis highlighted the fact that C2 and C3 were characterised by pathways related to viral response and interferon signalling (*Fig.S5E*), in agreement with the presence of IFNγ in innate-infiltrated regions. C4 also contained cytokine-related pathways. In addition, C0 and C1, which contained both innate-infiltrated and non-infiltrated cells, were characterised by pathways related to synapse formation and T-cell activation pathways (*Fig.S5E*), respectively, confirming that T-cell priming also occurred in regions that are not infiltrated by innate cells at this time point.

Taken altogether, our data confirms that CD8 T-cells present in innate-infiltrated and non-infiltrated regions can be activated by their TCR but experience different additional signals. The fact that CXCR3 and CD70 blockade abolishes T-cell expansion suggests that CD8 T-cell priming in non-infiltrated regions does not support proliferation.

### Treg segregation from initial priming sites provides a temporal window during which pathogen-specific T-cell priming and expansion is protected

We sought to understand why priming in innate-infiltrated supported T-cell expansion as opposed to priming in non-infiltrated regions. Because Tregs can directly impact T-cell priming^57,58^ and restrict T-cell expansion and contraction at specific, later times during infection^14,15,59^, we hypothesised that Tregs might be spatially regulated early during infection to allow for CD8 T-cell priming in innate-infiltrated regions and subsequent expansion.

To analyse where Tregs were located during CD8 T-cell priming, we stained for FoxP3 in spleen sections from mice infected by LM-OVA. Shortly after infection, Tregs distributed throughout the T-cell zone, as observed for steady-state, regardless of the presence of innate infiltration, highlighted by NK cells (*Fig.6A*). But over time, more than 60% of innate-infiltrated niches were characterised by Treg segregation from dense innate-infiltrated clusters (*Fig.S6A*). Using the density of NK cells to better delineate innate-infiltrated regions (*Fig.6B, S6B*), we found that Treg segregation started 18hrs post-infection (*Fig.6C*). Mixed regions, which contained both innate cells and Tregs, did not contain dense NK cell clusters (*Fig.S6B*), typically seen in innate-infiltrated regions devoid of Tregs (*Fig.6A-B*). Only a small proportion of Tregs expressed CXCR3 during LM-OVA infection (*Fig.S6C*), and CXCR3 expression on Tregs was not upregulated by TCR priming and/or short-term IFNγ treatment (*Fig.S6D*). We speculate that lack of CXCR3 upregulation following LM infection might provide a passive mechanism to segregate Tregs from innate-infiltrated regions.

**Figure 6.**
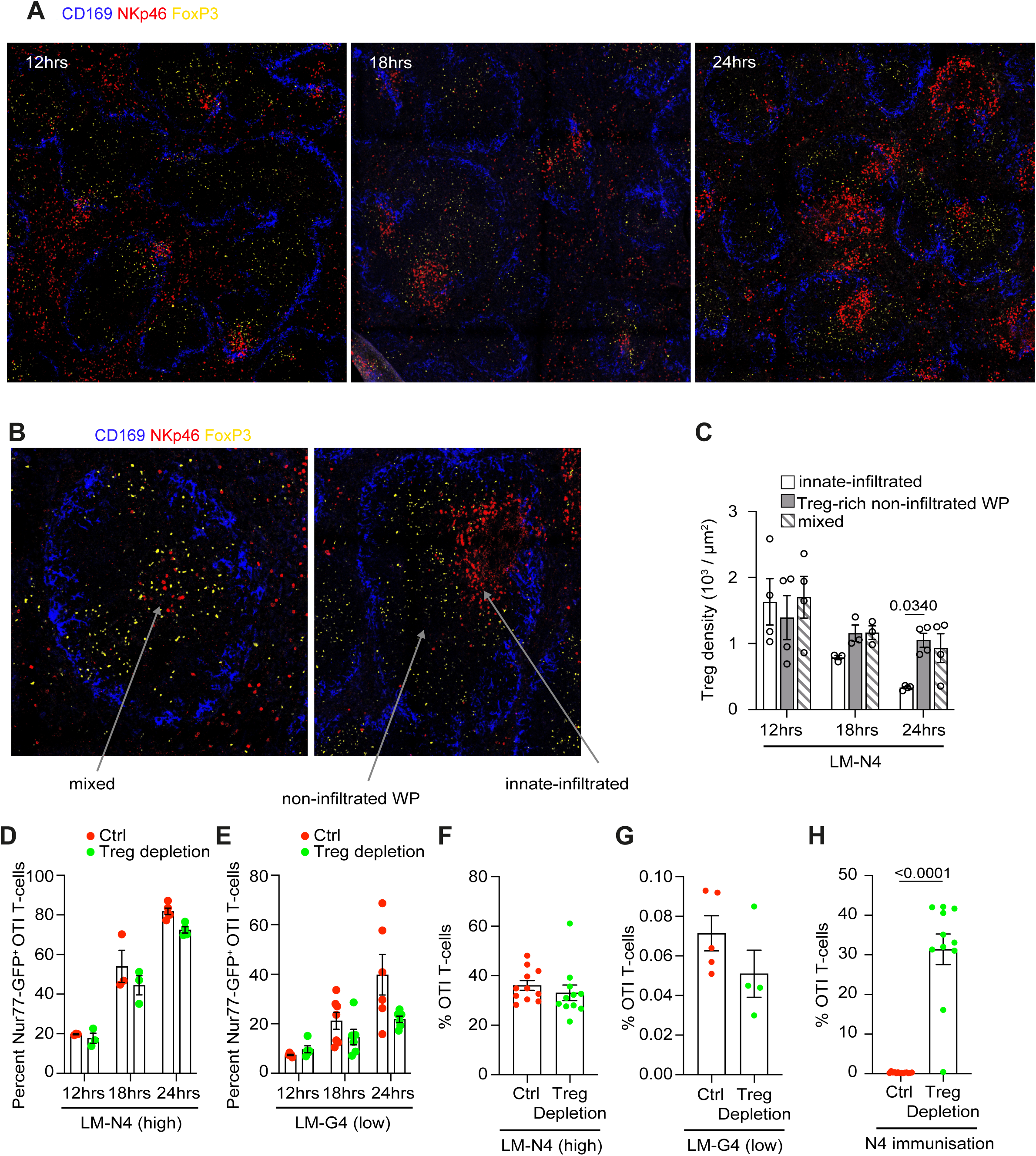
Tregs show spatial segregation away from innate-infiltrated regions and do not impact antigen-specific CD8 T-cell priming early during infection. **(A-C)** Spleen sections from mice infected by LM-OVA at the indicated time were stained for CD169 (blue), NKp46 (red) and FoxP3 (yellow) and analysed by microscopy. **A-** Representative images of Foxp3 localisation at the indicated times. **B-** Representative images of spleen section 24hrs after infection showing labelled innate-infiltrated, mixed, and non-infiltrated WP regions. **C-** Quantitation of FoxP3+ Treg densities within the indicated regions overtime (n=5-6 per timepoint). **(D-E)** WT and FoxP3-DTR mice were adoptively transferred with OTIxhCD2-DsRedxNur77-GFP T-cells, treated with DT and infected with LM-N4 or LM-G4. Graphs show the percentages of Nur77-GFP+ OTI T-cells at indicated timepoints following LM-N4 (**D**, n=3-5 per timepoint) and LM-G4 (**E**, n=10-13 per timepoint) infection at the indicated times post-infection. **(F-H)** WT and FoxP3-DTR mice were adoptively transferred with OTIxhCD2-DsRed cells, treated with DT and infected with LM-N4 **(F,H)** or LM-G4 **(G)** for 8 days. **(F-G)** Graphs show the frequencies of OTI T-cells after LM-N4 (**F**, n=11 per group) and LM-G4 (**G**, n=9-10 per group) infection. **H-** WT and Foxp3-DTR mice were transferred with OTI T-cells, treated with DT and immunised with the OVA peptide N4. Graph shows the frequency of OTI T-cells 5 days after infection (n=11 per group, 2 independent experiments).Two-way ANOVA with Turkey’s multiple correction test **(C)**, Welch’s t-tests with multiple comparisons correction using Holm-Šídák method **(D-H)**.

Treg segregation correlated with the kinetics of low-affinity T-cell priming, suggesting that, as a result of segregation, Tregs might not regulate initial CD8 T-cell priming. To address this, we used FoxP3-DTR mice to deplete Tregs following Diphtheria Toxin (DT) treatment^60^. DT treatment was performed in such a way that Tregs were depleted for the first 24hrs of infection (*Fig.S6E*). We then quantified OTI T-cell priming following infection with LM-N4 (high-affinity priming) or LM-G4 (low-affinity priming) using the reporter Nur77-GFP. Treg depletion did not result in earlier or increased priming of OTI T-cells, regardless of TCR affinity (*Fig.6D-E*), demonstrating that Tregs do not increase the threshold of TCR priming *in vivo* during LM infection. Similar data were obtained when we quantified the activation markers CD25 (*Fig.S6F-G*) and CD69 (*Fig.S6H-I*). Consistent with the fact that Treg depletion did not influence TCR priming, OTI T-cell expansion at the peak of the effector response was not affected by Treg depletion during priming, regardless of TCR affinity (*Fig.6F-G*). It was therefore unclear whether Tregs had any effect on pathogen-specific CD8 T-cell responses, as it did not impact their priming and expansion. Tregs have been shown to regulate T-cell fate during infection, and indeed, we found that Treg depletion during priming resulted in an increase in effector differentiation, based on the markers KLRG1 and CD127 (*Fig.S6J*). This indicated that Tregs specifically controlled pathogen-specific CD8 T-cell differentiation.

While Tregs depletion did not impact OTI T-cell expansion, we observed a general increase in CD8 T-cells at the peak of the effector response following Treg depletion (*Fig.S6K*), demonstrating that Tregs can limit overall CD8 T-cell expansion during infection and were therefore functional. It was then unclear why Tregs were unable to restrict OTI T-cell expansion. We therefore asked whether Tregs had the capacity to inhibit OTI T-cell expansion in other settings. We analysed OTI T-cell expansion following OVA peptide (N4) intravenous injection, an immunisation where antigen presentation was ubiquitous and did not result in inflammation (and thereby Treg segregation). Treg depletion during priming increased the number and frequency of effector OTI T-cells (*Fig.6H, S6L*), confirming the fact that Tregs have the capacity to inhibit OTI T-cell expansion when not segregated.

We conclude that Tregs are rapidly segregated from priming sites to facilitate high- and low-affinity T-cell priming. This initial priming is enough to drive pathogen-specific CD8 T-cell expansion, which we speculate becomes insensitive to Tregs. Altogether, these data strongly suggest that pathogen-specific CD8 T-cell priming is enabled by Treg segregation from initial inflammatory priming sites during infection.

## Discussion

Our study provides evidence that Listeria infection leads to a transient remodelling of the immune landscape characterised by the infiltration of NK cells and monocytes to the WP, coinciding with CD8 T-cell priming. In those innate-infiltrated regions, pathogen-specific T-cell priming is supported by CD70 expressed on cDC1. Importantly, the formation of innate-infiltrated regions is accompanied by the segregation of Tregs, essential to drive priming and expansion of pathogen-specific CD8 T-cells. Initial priming occurs in innate-infiltrated regions regardless of TCR affinity. This leads to the expression of CXCR3 in a TCR-affinity-dependent manner, which dictates the differential location of high- and low-affinity T-cells thereafter. As such, both the control of the TCR affinity threshold and priming of low-affinity T-cells rely on the transient formation of inflammatory niches, but the signals elicited by TCR affinity will subsequently affect their location and secondary signals received. Altogether, we propose that the transient formation of inflammatory niches, together with Treg segregation, is important to enable the priming of pathogen-specific CD8 T-cells, providing a window where both high- and low-affinity T-cells priming and downstream expansion is protected.

Innate infiltration coinciding with T-cell priming has been observed in other inflammatory conditions in lymph nodes^24,61^. This is driven by DCs that promote the expression of inflammatory chemokines and integrin ligands, necessary for efficient pathogen containment^61^. We therefore propose that the transient innate infiltration and Treg segregation we observe in the spleen during Listeria infection might constitute a general mechanism occurring during infection to improve pathogen-specific CD8 T-cell priming. Keeping innate infiltration limited is important for an effective immune response^61^, and we provide evidence that not only innate cells, but their downstream signalling cues are also confined. It was previously thought that cytokines permeate whole lymphoid organs during infection^62^. Our data suggests this is not necessarily the case, infection can drive the partitioning of specific cues, including localised expression of key costimulatory molecules such as CD70. This is to our knowledge one of the first reports demonstrating constrained costimulatory molecules expression in dedicated niches. It is also possible that other costimulatory molecules which are induced by inflammatory stimuli, such as OX40L^63^, are also expressed in a localised fashion.

Finally, we believe our study reframes the role of Tregs during infection. We provide clear evidence that Tregs are not implicated in pathogen-specific TCR priming and do not directly regulate the threshold of activation in this context. A seminal study provided evidence that depleting Tregs during priming decreases the avidity of the T-cell response^12^. Our study suggests that this might not be a direct role of Tregs on T-cell priming, but rather a consequence of the fact that low-affinity T-cells preferentially accumulate in non-infiltrated, Treg-rich regions before they have accumulated enough TCR stimuli to drive expansion. Previous studies have also suggested that Tregs are important at later stages, 3 to 7 days post-infection^14,15^ during which they regulate CD8 T-cell contraction and skew their fate towards memory. It was therefore not clear why Tregs affected CD8 T-cells during a secondary, but not primary phase of priming^15^. Our data, by providing evidence that Tregs are segregated from the initial priming site, explains why Tregs were tuning T-cell responses only at later times during infection. We propose that inflammation drives Treg segregation, due to their lack of CXCR3 expression. Interestingly, Tregs are able to express CXCR3 when subjected to long or chronic treatment with inflammatory cues, such as IFNγ-driven T-bet expression^17,64,65^ or CD40^66^. This suggests that, in chronic conditions, Tregs might have a direct negative impact on CD8 T-cell priming. Understanding the signals regulating Treg location in acute and chronic conditions might provide avenues to improve T-cell responses in vaccines and chronic conditions.

## Material and methods

### Mice

Mice were bred and housed in University of Oxford animal facilities, under specific pathogen-free conditions. Mice were routinely screened for the absence of pathogens and were kept in individually ventilated cages containing environmental enrichment at 20–24 °C, 45–65% humidity with a 12 h light/dark cycle. C57BL/6J (027), OTI (642) and CD45.1 (708) mice were purchased from Charles River. GREAT (IFNγ-reporter; 017580), Nur77-eGFP (018974), hUbi-GFP (GFP, 004353) and FoxP3-GFP-DTR (FoxP3-DTR, 016958) mice were purchased from Jackson Laboratory. OT3xTCR-α-/- mice^44^ were kindly gifted by Professor Zehn. hCD2-DsRed and hUbi-PA-GFP mice were kindly gifted by Professor Fiona Powrie at University of Oxford.

OTI mice were crossed with CD45.1 mice to generate congenically marked OTIxCD45.1 mice. OTI mice were crossed with hCD2-DsRed mice to generate OTIxhCD2-DsRed mice. OTIxhCD2-DsRed mice were crossed with Nur77-eGFP mice to generate OTIxhCD2-DsRedxNur77-eGFP mice. OT3xTCR-α-/- mice were crossed with hCD2-dsRed to generate OT3xTCR-α-/- hCD2-DsRed. hUbi-PA-GFP mice crossed with hCD2-DsRed mice to generate hUbi-PA-GFPxhCD2-DsRed mice.

Both male and female mice aged between 7 to 13 weeks were used for experiments, except for the bulk-RNA-seq experiments where only 8 week-old female mice were used.

All experiments involving mice were conducted in accordance with the United Kingdom Animal (Scientific Procedures) Act of 1986 and only procedures approved by the Home Office and Local Ethics Reviews Committee (University of Oxford) were performed, under the licences P4BEAEBB5 and PP3609558.

### Tissue harvesting and cell isolation

Spleens and lymph nodes were harvested from mice and mechanically homogenized by mashing though a 70μM cell strainer using a syringe plunger, and red blood cells were lysed in lysis buffer (155 mM NH_4_Cl, 12 mM NaHCO_3_ and 0.1 mM EDTA in ddH2O). Strained splenocytes were washed in R10 media (10% FBS, 1xPennicilin/Streptomycin/Glutamine [Gibco, 10378016], 50μM beta-mercaptoethanol [Gibco, 31350-010] in RPMI 1640 [Gibco, 21870076]) and counted.

In some experiments spleens were dilacerated using scalpels and incubated with 100μg/ml Liberase™ TM (Merck, 5401119001) and 10μg/ml DNAse I (PanReac AppliChem, A3778,0100) for enzymatic digestion. The digested tissues were then homogenized though a 70μM cell strainer and treated with red blood cell lysis buffer as previous steps.

CD8 T-cell isolation was conducted using a mouse CD8 T-cell isolation kit (Miltenyi Biotec, 130-104-075), in accordance with the kit protocol.

### In vitro stimulation experiments

Isolated CD8 OTI T-cells were co-cultured with C57Bl6 splenocytes in a 2:1 ratio in 96-well plate flat-bottom plates. The splenocytes were pre-incubated with recombinant mouse IFN-γ (BioLegend, 575306) at 10ng/ml for 2hr prior to the addition of CD8^+^ OTI cells. Altered OVA peptide ligands, SIINFEKL (N4) and SIIGFEKL (G4) (Proteogenix) were added to the co-culture at 50ng/ml. At specified timepoints, cells were harvested for flow cytometry.

### Infection and in *vivo* treatments

LM-OVA^67^, LM-N4^31^ and LM-G4^31^ (provided by Prof. Zehn) were expanded in Brain Heart Infusion (BHI) broth (Sigma, 53286-100G) at 37◦C and diluted in PBS once the exponential growth phase was reached (OD600 measurements between 0.075-0.1). Mice were infected via tail vein intravenous injection (8×10^3^ CFU for 8-day experiments and 2×10^4^ or 5×10^4^ CFU for 24hr experiments). LM suspensions were plated on BHI agar plates (Sigma, 70138-500G) for colony counting following overnight culture at 37◦C.

In some experiments, OTI splenocytes (1×10^7^ cells for 24hr experiments) or isolated OTI T-cells (3×10^6^ cells for 8-day experiments and 5×10^4^ cells for 8-day experiments) were resuspended in PBS and transferred into recipient mice via intravenous injection 24hr prior to infection, unless otherwise stated.

In some experiments, isolated CD8 T-cells were resuspended in PBS containing 5μM eBioscience™ CFSE (Invitrogen, 65-0850-85) or 1μM CellTrace™ Far Red dye (Invitrogen, C34564) diluted in PBS, for 20min at room temperature in darkness. Cells were washed twice in complete R10 media and resuspended in PBS prior to cell transfer.

For CXCR3 blockade, mice were intraperitoneally administered with 300μg anti-CXCR3 (BioLegend, 126537) or isotype control antibody (BioLegend, 400959) 6hr before infection, unless otherwise stated.

For CD70 blockade, mice were intraperitoneally administered 300μg anti-CD70 antibody (BioXCell, BE0022) or isotype control antibody (BioXcell, BE0290) 6hr before infection, either alone or in combination with CXCR3 blockade.

For Treg depletion experiments, FoxP3-DTR mice were intraperitoneally administered 500ng diphtheria toxin (Merck, 322326) on two consecutive days.

For OVA immunisation, mice were administered 200ug SIINFEKL (N4) peptide (ProteoGenix) via tail vein intravenous injection.

For the imaging of cytokines, mice were intraperitoneally administered with 250µg Brefeldin A (BFA) (Cayman Chemicals) 6hr prior to tissue harvest.

### Immunofluorescence imaging

Freshly harvested spleens were cut in halves and fixed in paraformaldehyde-lysine-periodate solution (1% PFA, 75 mM L-lysine [Sigma, L5751], 10 mM sodium m-periodate [Thermo Scientific™, 20504], diluted in phosphate buffer containing 0.2M NaH_2_PO_4_ [Sigma, S3139] and 0.2M Na_2_HPO_4_ [Sigma, S0876], adjusted to pH 7.4) for 16-24hr at 4°C with gentle agitation. After washing in 1XPBS 1 hr at 4 °C with gentle agitation, the spleens were dehydrated in 30% sucrose for 24-48hr, embedded in Optimal Cutting Temperature Compound (VWR, 361603E), and frozen in dry ice bath with methanol. The frozen spleens were stored at −80 °C prior to cryosectioning at 10 μm thickness using a Leica CM1900UV cryostat onto Superfrost Plus microscope slides (VWR, 631-0108).

Cryosections were washed with PBS, permeabilized and blocked with imaging buffer (2% FBS, 0.01% sodium azide,0.3% Triton X-100 (Thermo Scientific™, A16046.AE) in PBS) containing TruStain FcX^TM^ (BioLegend, 101320) and species-specific serum depending on the fluorophore-conjugated antibodies for 2-4hr at room temperature. For NKp46 staining, the blocked sections were incubated with unconjugated anti-NKp46 overnight at 4°C. For the secondary detection of anti-NKp46, the sections were washed three times with imaging buffer and stained with fluorophore-conjugated secondary antibody for 2hr at room temperature. The sections were washed and blocked again for 1-2hr before incubation with fluorophore-conjugated antibodies overnight at 4°C. After final washing, the sections were mounted with Fluoromount G (Southern Biotech, 0100-01) and covered with glass coverslips on top. For 33D1 staining, imaging buffer made with PBS containing Ca^2+^ and Mg^2+^ was used. Images were acquired at 10X magnification using either ZEISS LSM 980 Airyscan confocal microscope or ZEISS Axioscan7 Slide Scanner with Zen software (Carl Zeiss).

### Quantification of inflammatory regions, cell infiltration, and Nur77-GFP+ OTI cells

All immunofluorescence images were processed and analysed using the ImageJ software (V2.16.0). Prior to analysis, images were quality-inspected for major tears or folds. Images with Z-stacks were subject to average intensity Z-projection and were presented as composite images. Regions of interest (ROIs) were manually delineated on each tissue section based on CD169 staining for white pulps and NK cells within white pulps for innate-infiltrated regions. For cell counting, the corresponding fluorescence channels were manually thresholded to distinguish cell objects from background, and watershed separation was applied to separate any adjoining multiplets when appropriate. Cell objects were quantified using the “Analyse Particles” function with a minimum size of 10μm^2^ for NK cells and CD8 T cells, and 7μm^2^ for FoxP3+ cells. ROIs containing clustered cell objects that cannot be resolved were excluded from further analysis due to the inaccuracy of cell numeration. NK-infiltrated regions were defined as ROIs with a minimum cell density of 1 cell per 10^3^μm^2^. For the quantification of Nur77-GFP+ OTI cells, the respective fluorescence channels were individually thresholded and merged. The merged images were then converted to RGB images for colour thresholding of double positive cells. For RGB images converted from merged channels that are red and green, objects displaying Hue values between 20-50 were counted as double-positive.

### Flow Cytometry

Cells were stained in 96-well V-bottom plates (Corning, 3894). Fc-receptor blocking and live/dead staining were performed simultaneously by incubating cells in Zombie NIR dye (BioLegend, 423105) and TruStain FcX^TM^ (BioLegend, 101320) in PBS for 15min on ice. For extracellular staining, cells were incubated in fluorophore-conjugated antibodies diluted in flow cytometry buffer (2% FBS, 2.5mM EDTA, 0.01% sodium azide in PBS) for 30min on ice or overnight at 4°C. For 33D1 staining, flow cytometry buffer made with PBS containing Ca^2+^ and Mg^2+^ was used. For tetramer staining, cells were stained with AF647- or BV421-conjugated N4-MHC I tetramers (NIH Tetramer Core Facility) in flow cytometry buffer for 30min at room temperature following extracellular staining. Unless otherwise stated, cells were fixed in 4% paraformaldehyde (Thermo Scientific™, 11586711) in PBS for 20 minutes on ice. For intracellular staining, cells were fixed and permeabilized with either BD Cytofix/Cytoperm^TM^ kit (BD Biosciences, 554714) for cytosolic targets or True-Nuclear™ Transcription Factor Buffer Set (BioLegend, 424401) for transcription factors, in accordance with the kit protocols. Fixed and permeabilized cells were incubated with fluorophore-conjugated antibodies diluted in kit-specific permeabilization buffers for 30min on ice. All samples were resuspended in flow cytometry buffer prior to data collection. For some experiments, CountBright™ Absolute counting beads (ThermoFisher, C36950) were also added. Data were recorded on BD LSRII or FortessaX20 using BD FACSDiva software (V8.0) and analysed on FlowJoTM software (V10.10.0) or recorded on Cytek 4-laser Aurora using Cytek SpectroFlo software and analysed using Omiq (http://www.omiq.ai/).

### Two-photon microscopy of explanted spleens

Mice were transferred with a 1:1 mix of 6×10^6^ OTI GFP and OT3 cells, 6×10^6^ OTI or 6×10^6^ OT3 24hrs prior infection with 20.000 CFU of LM-OVA or injection with PBS. Spleens were explanted 24hrs p.i, 300-400 sections were sliced with a compresstome (VF210-OZ) and stained for minimum 30 min with CD11b BV421. Innate-infiltrated and non-infiltrated regions were identified by the presence/absence of CD11b+ clusters, respectively. Sections were then immobilized in coverslips before acquisition.

Images were acquired with Zeiss LSM 880 microscope with a W Plan-Apochromat 20 x 1.0 DIC D = 0 vis-IR M27 75 mm objective and two Mai Tai lasers with a range of excitation of 690-849 and 850-990 nm. Detectors used include a nose piece GaAsp1 with filters allowing detection of 425/60nm for blue dyes and CFP, a nose piece GaAsp2 with 500-550 nm filters for detection of green dyes including GFP, YFP and CFSE, and a BIG2 GaAsp detector with filters range of 590-610 nm for red dyes and red fluorescent proteins. Videos were acquired with Zen software (Carl Zeiss, Inc.) Cells stained with CD11b-BV421 were exited with 810nm and all fluorescent proteins were excited with 930nm. Time Series were acquired in planes of 4 to 5µm z-spacing spanning a depth of 24-64µm every 30s for 35-40 min.

Time series videos were analysed with Imaris Bitplane (V9.7.2). Cells were quantified and tracked using “spots” feature and considered arrested when they reduce the speed (change of event) below 2.5 µm/min for at least 2 consecutive tracks. The coefficient of arrest was then calculated as the percentage of the full length of the track that each cell remained arrested.

### Photoactivation experiments

hUbi-PA-GFP hCD2-DsRed mice were infected with 20.000 CFU of LM or injected with PBS. Spleens were collected 24hrs p.i, sliced and stain for CD11b-BV421 as described above. Innate-infiltrated and non-infiltrated regions in individual sections were identified by CD11b stain and DsRed+ T-cells from each region were photoactivated by irradiation with Mai Tai II laser (wavelength 840nm; laser power intensity 60-80%; acquisition speed 6s with 2 iterations). For an efficient photoactivation, z-depth was kept below 40 µm. Successful photoactivation was observed by GFP fluorescent at 940nm. Non-photoactivated hUbi-PA-GFP hCD2-DsRed mice sections were used as negative control. An average of 6 niches per section and 3 sections/experiment were photoactivated depending on the quality and size of the section. Spleens from two mice were processed in each experiment.

LM-infected and naive spleen sections were collected in a 6-well plate and mechanically smashed into single cell suspensions. Splenocytes were stained with the following antibodies: CD8a BV711 (clone 53-6.7), CD4 AF647 (clone RM4-5) and Fixable Viability Dye eFluorTM 780 (eBioscience) and GFP+, photo-activated CD8 T-cells were sorted. One-thousand cells were collected in 6µl of RNA-lysing buffer (NEBNext Cell Lsysis Buffer, New England Biolabs Inc.) per experiment. Samples were collected in triplicate out of 6 independent experiments and processed for ultra-low RNA bulk sequencing by the Oxford Genomic Centre, Oxford, UK.

#### Bulk RNA-seq analysis

Bulk RNA reads were mapped using Hisat2 (2.2.1) and GRCm39 (v111) from Ensembl. Genes were filtered for minimal expression across multiple samples and signature cell-type genes (CD4, CD19, ITGAX, KLRB1, NCR1) were examined to check for potential contaminating cell-types. Samples found to include CD4 T cells, B cells, Macrophages or NK cells were excluded from further analysis. Differential expression analyses were performed using edgeR’s voomLmfit (edgeR_3.34.1), blocking on biological replicate. Three separate differential analyses were performed to compare naive versus innate-infiltrated, naive vs non-infiltrated, and innate-infiltrated vs non-infiltrated. Gene set enrichment analysis was performed with gseGO (clusterprofiler).

#### Single Cell RNA-seq analysis

Sequence reads were mapped using CellRanger multi (version 6.0.0) with the 10x mouse reference transcriptome (version 2020-A). Datasets were analysed using Seurat version 4.0.5^68^. Samples were merged on a single Seurat object. We filtered out cells having less than 200 and more than 5,000 detected genes, cells in which mitochondrial protein-coding genes represented more than 10% of UMI and ribosomal protein-coding genes represented more than 50% of UMI. Data was processed using the standard Seurat (V5.4.0)^69^ workfow (NormalizeData, FindVariableFeature, ScaleData) and variation associated with mitochondrial UMI percentage was regressed out. Principal components were calculated using the top 3,000 variable features. These genes were used as input for principal component analysis (PCA), and 30 PCs was used for UMAP and clustering. Clusters corresponding to memory T cells (based on CD44, Eomes and Tbx21 expression) were removed and clustering was re-run. Final clustering was performed with the Louvain algorithm (n=30 PCs, resolution=0.3). Significant differentially expressed genes between clusters were identified using the “FindAllMarkers” function, Wilcoxon test and selecting markers expressed in at least 25% of cells. Pathway analysis was performed with g:Profiler^70^. Gene signature of cells originating from naïve mice, innate-infiltrate regions or non-infiltrated regions was extracted from bulk-RNAseq (*Dataset 3-4*), using SingleR^71^ for cell annotation.

### Statistical Analysis

Two-tailed Welch’s *t*-tests were used for statistical comparisons between two biologically unmatched experimental groups. Two-tailed paired *t*-tests were used for comparisons between two biologically-matched groups. ANOVA tests were used for comparisons across three or more experimental groups. Data were considered statistically significant when p < 0.05. Data are presented as mean ± SEM unless specified. All statistical analyses were performed using Graphpad (V10.0.3, Prism software).

## Data availability

The mouse bulk RNAseq data generated in this study is been deposited the GEO database. Datasets reused in this study: <u>GSE244203</u>^35^. All R scripts used for performing the bulk-RNAseq analysis are available on Github: https://github.com/muir-a/Muir-Gerard-SplenicCompartments

## Author Contributions

**A.L.C.:** Conceptualization, Investigation, Formal analysis, Visualization, Writing - Original Draft; **H.C.:** Conceptualization, Investigation, Formal analysis, Visualization, Validation, Writing - Original Draft; **G.J.M.:** Investigation, Validation, Writing - Review & Editing; **A.M.:** Formal analysis, Visualization, Writing - Review & Editing; **M.B.:** Investigation, Writing - Review & Editing LFKH: Investigation, Writing - Review & Editing; **A.C.R.:** Formal analysis, Visualization, Writing - Review & Editing, Supervision, Funding acquisition; **A.G.:** Conceptualization, Formal analysis, Visualization, Writing - Original Draft, Supervision, Project administration, Funding acquisition

## Acknowledgments

We would like to thank the Wellcome Trust Centre for Human Genetics for the generation of the sequencing data, J. Webber for the assistance with cell sorting, the dynamic platform and microscopy facility at the Kennedy Institute, and Dietmar Zehn and Fiona Powrie for mice and reagents. We would also like to thank Tal Arnon and Ed Roberts for critical reading of the manuscript. This work was supported by the UKRI (BBSRC BB/R015651/1 to A.G.), the John Fell Funds (0006162 to A.G.), Cancer Research UK (CR-UK) (DRCPFA-Nov23/100006 to A.G.); the Kennedy Trust for Rheumatology Research (KENN151607 and KENN202112 to A.G.), and Kennedy Trust and MRC studentship (to H.C.). A.M. was supported by an Institute Strategic Programme Grant to the Babraham Institute (BBS/E/B/000C0545), and A.C.R. by a UKRI Medical Research Council Career Development Award (MR/W016303/1).

---

**Figure S1.**
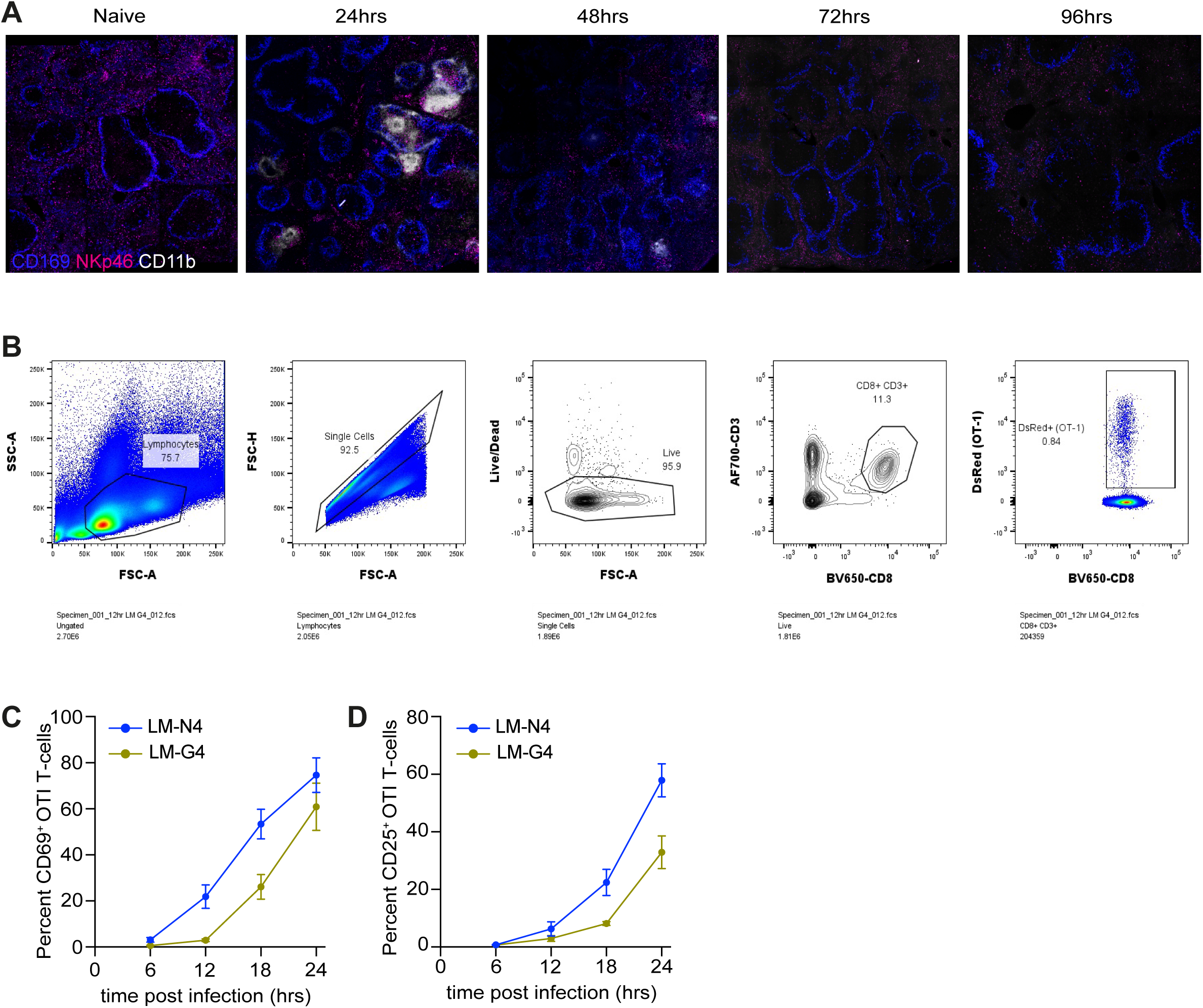
CD8 T-cell priming is spatiotemporally linked to inflammatory innate cell infiltration following LM infection. **A-** Representative images of spleens from mice infected with LM-OVA and harvested at indicated time points post-infection. CD169 (blue), CD11b (white), NKp46 (magenta). **(B-D)** WT mice were adoptively transferred with OTIxhCD2-DsRedxNur77-GFP T-cells and infected with LM-N4 (blue) or LM-G4 (yellow). Spleens were harvested at indicated time points. **B-** Gating example for the analysis of OTI T-cells. The percentages of CD69+ **(C)** and CD25+ **(D)** OTI T-cells was identified by flow cytometry (n=4 for 6hr, n=12 for 12hr, n=16 for 18hr, n=17 for 24hr).

**Figure S2.**
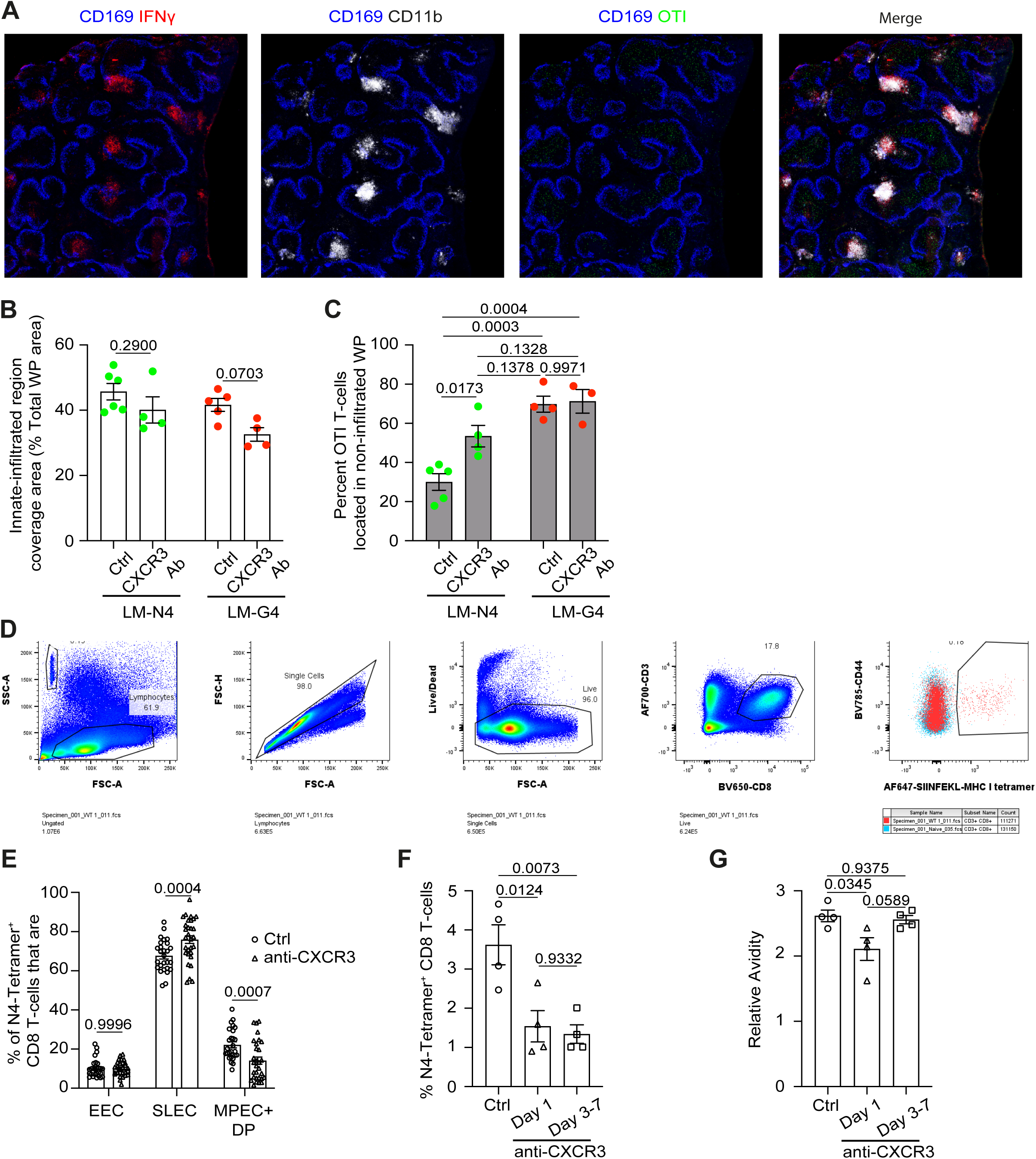
CXCR3 expression on high-affinity CD8 T-cells allows their retention in innate-infiltrated regions and promotes expansion. **A-** Mice were adoptively transferred with OTIxGFP T-cells, infected with LM-OVA and treated with BFA 6hrs prior to spleen harvest. Representative images of spleen section stained for CD169 (blue), IFNy (red), CD11b (white), OTI (green) 24hrs after infection. **(B-C)** Mice were adoptively transferred with OTIxhCD2-DsRed T-cells, infected with LM-N4 or LM-G4, and treated with either anti-CXCR3 or isotype control 6hr prior to infection. After 24hrs, spleen sections were stained for CD169 (blue) and NKp46 (magenta) (n=3 per group). **B-** Area coverage of innate-infiltrated regions as percentages of total white pulp areas on stained spleen sections. **C-** Percentage of OTI T-cells located within non-infiltrated regions. **(D-F)** Mice were infected with LM-OVA and treated with anti-CXCR3 or isotype control either once 6hr before infection **(E)** or on day 3, 5, and 7 post-infections **(F-G)**. Spleens were harvested 8 days post-infection and analysed flow cytometry (n=4-5 per group). **D-** Gating example for the analysis of OVA-specific CD8 T-cells. **E-** Relative abundance of early effector cells (EEC, CD44+ KLRG1-CD127-), short-lived effector cells (SLEC, CD44+ KLRG1+ CD127-), and memory precursor effector cells (MPEC, CD44+ KLRG1- CD127+) among N4-tetramer+ CD8 T-cells. **F-** Frequencies of N4-tetramer+ CD8 T-cells. **G-** Relative avidity of N4-tetramer+ CD8 T-cells normalised to isotype control. Two-way ANOVA with Turkey’s multiple comparison test (B-c, E), Welch and Brown-Forsythe one-way ANOVA (F-G).

**Figure S3.**
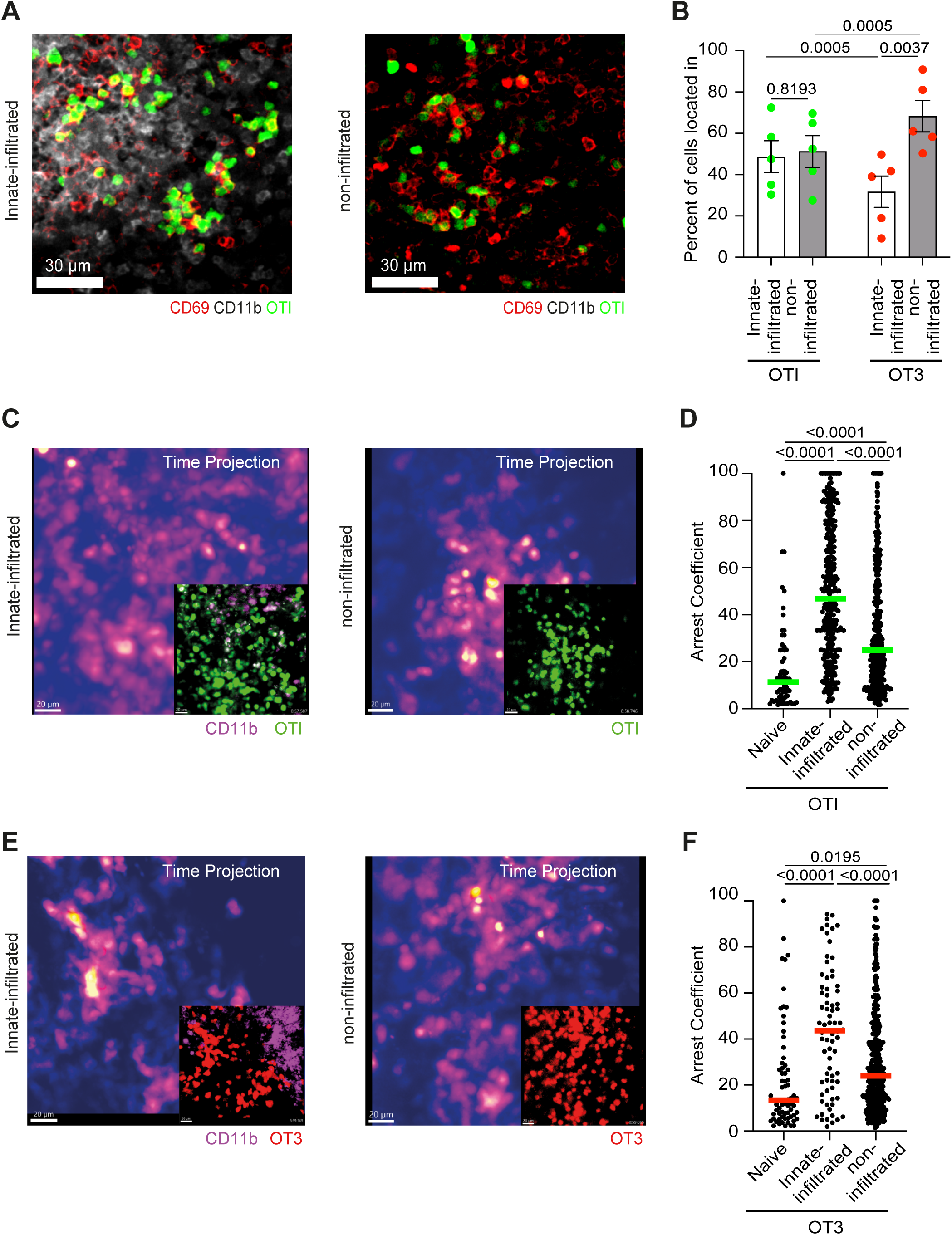
CD8 T-cells encounter antigens in both innate-infiltrated and non-infiltrated regions, regardless of TCR affinity. Mice were adoptively transferred with OTIxGFP **(A,C-D)**, OT3xhCD2-DsRed **(E-F)** cells separately or admixed **(B)** and infected with LM-OVA. After 24hrs, live spleen sections were labelled with Cd11b and imaged by 2-photon microscopy. Data is representative from 2 independent experiments with at least 3 spleens per condition. **A-** Image of innate-infiltrated (left) and non-infiltrated (right) region of a WP showing clustering of OTI T-cells (green), CD11b (white) and CD69 (red). **B-** Percentages of OTI (green) and OT3 (red) T-cells located within the innate-infiltrated (white) or non-infiltrated (grey) regions. **(C-F)** Pseudocolored time projection of a 30-minute timelapse **(C,E)** and arrest coefficient **(D,F)** showing the spatial persistence of OT-I **(C-D)** and OT3 **(E-F)** T-cells in innate-infiltrated and non-infiltrated regions. **(D, F)** Each dot is a cell, bar indicates Median.

**Figure S4.**
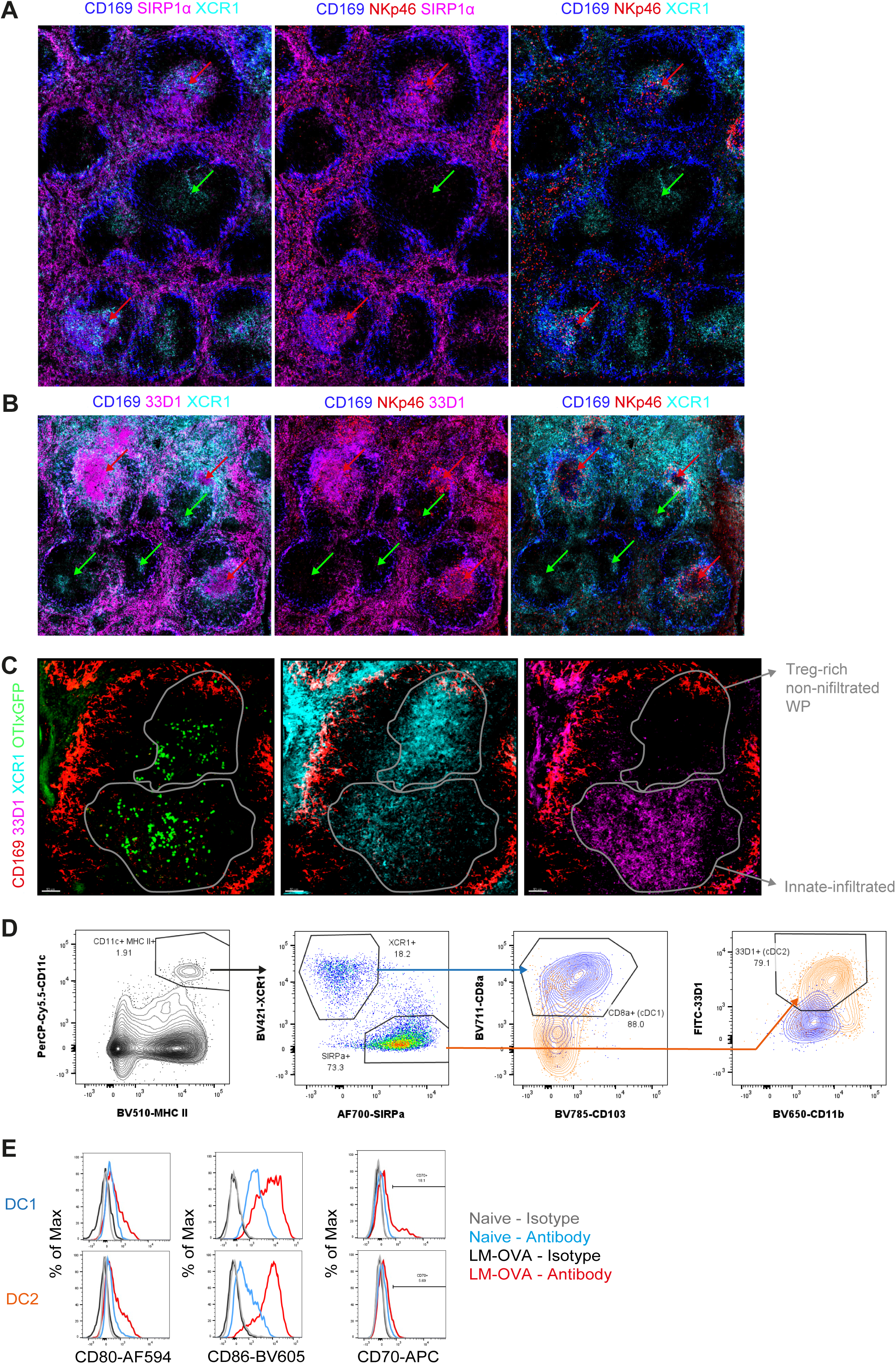
Inflammatory innate infiltration is accompanied by DC spatial redistribution and co-stimulatory molecule expression. **(A-B)** Representative image of spleen sections from mice infected with LM-OVA for 24hrs were stained for CD169 (blue), XCR1 (cyan), NKp46 (red) and SIRPa (**A**, magenta), or 33D1 (**B**, magenta). **C-** Mice were adoptively transferred with OTIxGFP cells and infected with LM-OVA. Representative image of spleen sections stained for CD169 (red), 33D1 (magenta), XCR1 (cyan), OTI (green) 24hrs post-infection. **(D-E)** Splenocytes from mice infected with LM-OVA for 24hrs were analysed by flow cytometry. **D-** Representative gating strategy for splenic cDC1 and cDC2 subsets. **E-** Representative histograms showing CD80, CD86, and CD70 expression in cDC1 and cDC2.

**Figure S5.**
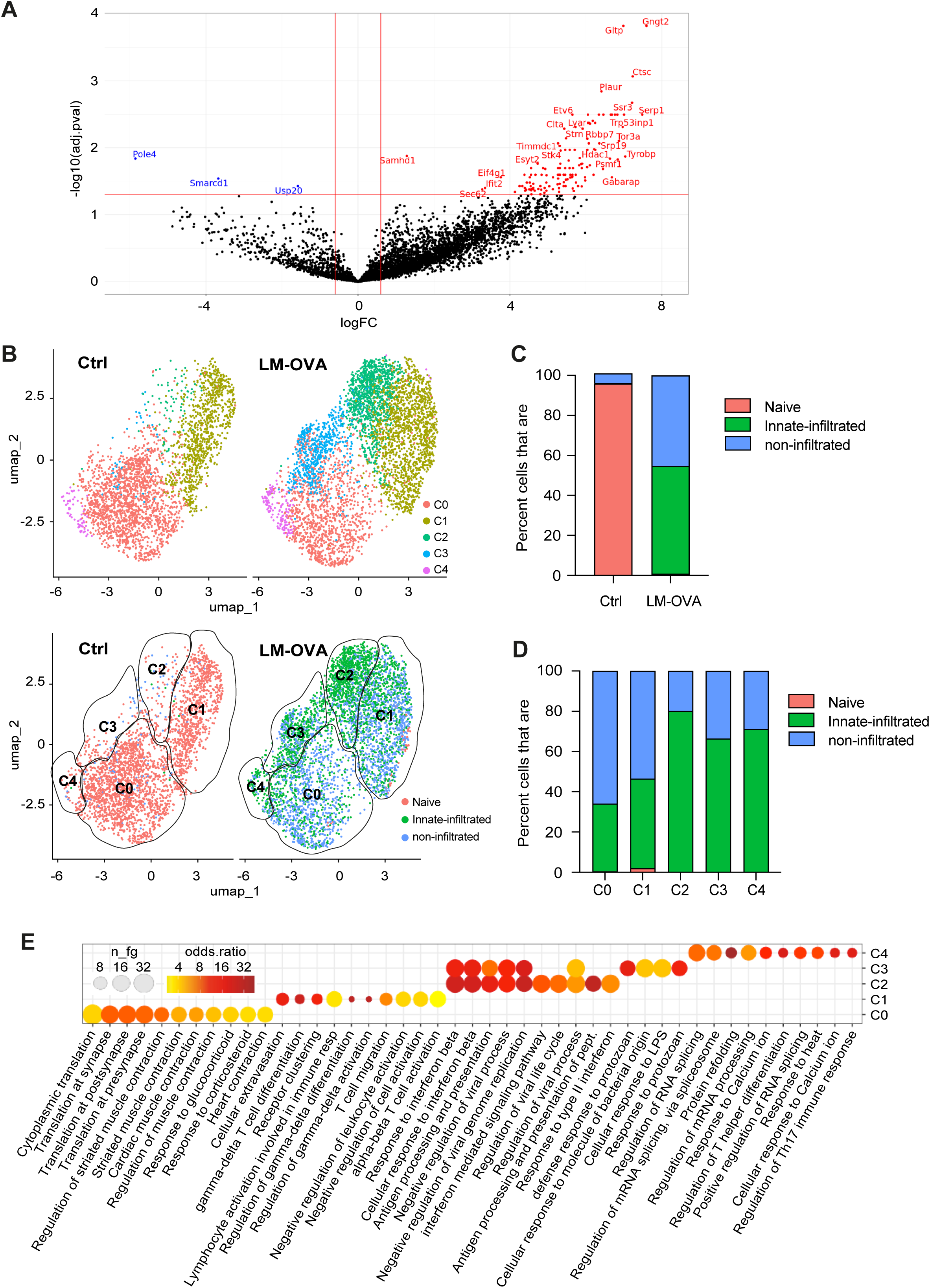
Transcriptomic signature associated with innate infiltration is only present in a subset of TCR-primed CD8 T-cells. **A-h**CD2-DsRedxPA-GFP mice were infected with LM-OVA. After 24hrs, spleens were explanted, sectioned and stained for Cd11b and Ly6C. Innate-infiltrated and non-infiltrated regions were highlighted by 2-photon light to activate PA-GFP. PA-GFP+ cells were then sorted and subjected to bulk-RNAseq. Volcanoplot of differentially expressed genes significantly up-regulated (red) and down-regulated (blue) in innate-infiltrated niches compared to non-infiltrated niches. **(B-E)** Analysis of scRNAseq of non-memory CD8 T-cells from spleen of control (ctrl) or LM-OVA-infected mice for 24hrs. Cells were clustered and their putative location was assigned according to their expression of Naive, innate-infiltrated or non-infiltrated signature from Fig.5 (n=2931 for ctrl, 4522 for LM-OVA). **B-** UMAP visualization of cells according to clusters (top) or gene signature (bottom; clusters highlighted with black contour). **C-** The bar plot shows proportion of cells from each niche within ctrl or LM-OVA samples. **D-** The bar plot shows proportion of cells from each niche within each cluster, focussing on the LM-OVA condition. **E-** Dot plot heatmap showing GO biological pathways significantly enriched across each cluster.

**Figure S6.**
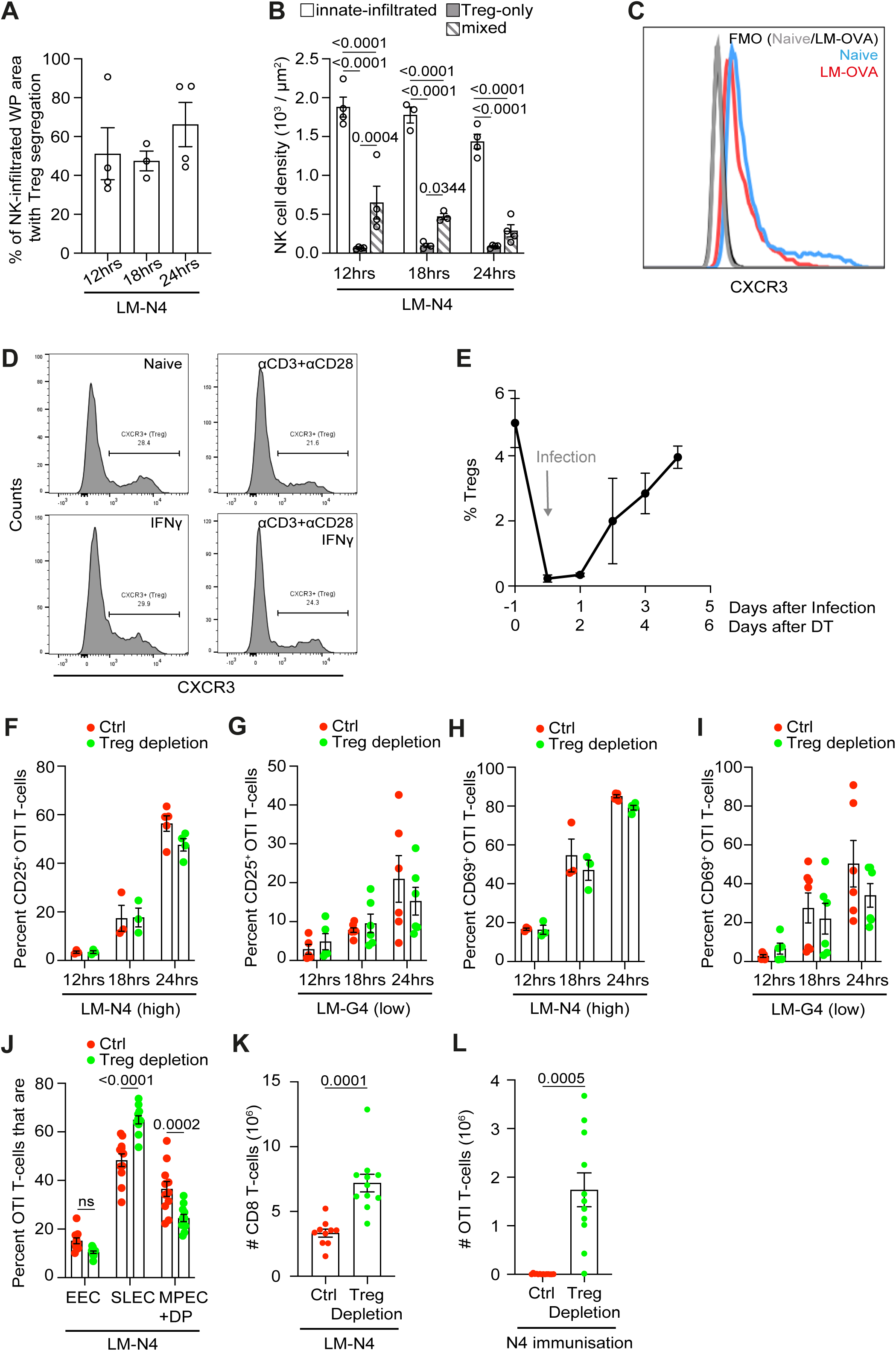
Tregs are spatially segregated away from innate-infiltrated regions and do not affect antigen-specific CD8 T-cell priming early during infection. **(A-B)** Spleen sections from LM-OVA infected mice were stained for CD169 (blue), NKp46 (red) and FoxP3 (yellow) (n=5-6 per timepoint) at the indicated times and analysed by microscopy. **A-** Relative abundance of NK-infiltrated regions exhibiting Treg segregation, shown as percentages of total white pulp areas. **B-** Quantitation of NK cell densities within different WP regions. **C-** Representative histogram showing CXCR3 expression on splenic Tregs from naive mouse (blue) or mouse infected for 24hrs with LM-OVA (red). **D-** Splenocytes were isolated and stimulated in vitro as indicated for 24hrs. Histograms show the expression of CXCR3 in Tregs. **E-** FoxP3-DTR mice were treated with DT as indicated and infected with LM-OVA. Graph shows the frequency of Tregs in peripheral blood over time. **(F-I)** WT and FoxP3-DTR mice were adoptively transferred with OTIxhCD2-DsRedxNur77-GFP T-cells, treated with DT and infected with LM-N4 or LM-G4. Graphs show the frequency of CD25+ **(F-G)** or CD69+ **(H-I)** OTI T-cells with LM-N4 (**F, H**, n=3-5 per timepoint) and LM-G4 (**G, I**, n=10-13 per timepoint) at the indicated times. **J-** WT and FoxP3-DTR mice were adoptively transferred with OTIxhCD2-DsRed cells, treated with DT and infected with LM-N4 for 8 days. Relative abundance of early effector cells (EEC, CD44+ KLRG1-CD127-), short-lived effector cells (SLEC, CD44+ KLRG1+ CD127-), memory precursor cells (MPEC, CD44+ KLRG1- CD127+) and double positive cells (DP, CD44+ KLRG1+ CD127+) OTI T-cells. **K-** WT and Foxp3-DTR mice were treated with DT and infected with LM-OVA. Graphs show the absolute number of total splenic CD8 T-cells 8 days after infection (n=10-11 per group). **L-** WT and Foxp3-DTR mice were transferred with OTI T-cells, treated with DT and immunised with the OVA peptide N4. Graph shows absolute numbers of OTI T-cells 5 days after infection (n=11 per group, 2 independent experiments). Two-way ANOVA with Turkey’s multiple correction test **(B, F-J)**, Welch’s t-tests **(K-L)**.

